# ATP13A2 Loss of Function-Driven Polyamine Dysregulation Induces SAM Depletion and Epigenetic Astrocyte Toxicity

**DOI:** 10.64898/2026.04.02.716164

**Authors:** Elena Coccia, Gustavo Morrone Parfitt, Sundas Ijaz, Aisha Sati, Julia Gesner, Andrea Perez Arevalo, Jennifer Strong, Anna Bright, Soha Sohail, Thea Meimoun, Tim Ahfeldt, Peter Vangheluwe, Joel Blanchard

## Abstract

Aging is the strongest risk factor for neurodegeneration, yet how the human brain ages remains poorly understood. Loss-of-function (LOF) variants in *ATP13A2* cause severe juvenile-onset Parkinson’s disease, providing a window into the mechanisms that accelerate age-related neurodegeneration. ATP13A2-LOF causes lysosomal polyamine sequestration, but how this promotes pathogenesis remains unclear. We discovered that ATP13A2-LOF depletes cytosolic polyamines in astrocytes, triggering compensatory upregulation of *de novo* polyamine biosynthesis, which diverts S-adenosyl methionine (SAM) from DNA and histone methylation, leading to increased chromatin accessibility and epigenetic reprogramming of astrocytes into a neuroinflammatory state that releases neurotoxic cytokines that promote dopaminergic neuron death. In ATP13A2 knockout mice and human models, we find that genetic and pharmacological inhibition of SAM utilization in polyamine biosynthesis prevents astrocytic epigenetic reprogramming and promotes dopaminergic neuron survival. These findings reveal a direct link between polyamine metabolism, epigenetic dysfunction, and neurotoxic inflammation, uncovering new therapeutic opportunities in Parkinson’s disease.

## Introduction

Parkinson’s Disease (PD) is a debilitating neurodegenerative disease defined by the selective vulnerability and loss of dopaminergic neurons (DN), resulting in a spectrum of motor and non-motor symptoms^1^. Although decades of research have identified key pathogenic pathways, including impaired protein homeostasis, neuroinflammation, lysosomal dysfunction, and mitochondrial impairment^2,3^, how these processes converge at the cellular level to promote disease onset and progression is incompletely understood.

Genetic forms of PD provide a powerful framework to address this gap by linking defined molecular perturbations to disease mechanisms. Approximately 5% of PD cases are associated with rare genetic variants that cause early-onset familial disease and often exhibit severe clinical presentations. Among these, loss-of-function (LOF) mutations in the lysosomal protein ATP13A2 (*PARK9*) cause one of the earliest and most aggressive forms of PD with onset in adolescence and early adulthood^4–6^. Converging genetic and mechanistic studies have positioned lysosomal dysfunction as a central mechanism in PD pathogenesis, largely thought to be driven by degradation and clearance of neurotoxic proteins such as α-synuclein^7–9^. However, ATP13A2 encodes a lysosomal P-type ATPase that exports polyamines (PAs) from the lysosomal lumen into the cytosol, thereby regulating their intracellular compartmentalization of PAs. PAs are essential metabolites that support diverse cellular processes, including autophagy, resistance to oxidative stress, mitochondrial function, and receptor signaling^10–14^. The highly penetrant juvenile onset of Parkinsonian symptoms caused by mutations in *ATP13A2,* therefore highlight the potential importance of PA bioavailability in specific cell compartments, rather than abundance as a determinant of PD pathogenesis. Consistent with this, alterations in PA metabolism were linked to PD and neurodegeneration, even before ATP13A2 was identified as a lysosomal PA transporter in 2020^15–19^. Beyond neurodegeneration, PA homeostasis has been strongly associated with aging and organismal longevity: endogenous levels of PAs decline with age, pointing to their role in age-related cellular processes^20,21^, and supplementation with spermidine extends lifespan in yeast, worms, flies, and mice^22,23^. Together, these observations position ATP13A2 at the intersection of lysosomal function, metabolic regulation, and age-related neurodegeneration. However, how lysosomal PA dysregulation leads to pathogenic cellular states remains unknown.

Previous studies in cell culture models have shown that ATP13A2 dysfunction impairs lysosomal function and disrupts α-synuclein handling^5,13,24–26^. Yet this framework alone does not fully explain disease progression: ATP13A2-associated PD does not consistently feature prominent α-synuclein pathology, and animal models exhibit early gliosis that precedes neuronal loss and motor phenotypes^27–30^. These observations point to a particular sensitivity of glial cells rather than neurons alone to PAs dysregulation. Accumulating evidence further positions astrocytes as drivers of neurodegeneration, whereby loss of homeostatic astrocytic functions and acquisition of neurotoxic, pro-inflammatory states contribute directly to neurodegeneration^31–33^. Astrocytes are also key regulators of brain PA metabolism: PAs enter the brain via astrocytic transporters at the blood-brain barrier, are stored within astrocytes, and are released as neuromodulators to support neuronal health^34–36^.

Emerging work has highlighted epigenetic regulation as a critical determinant of glial identity and inflammatory potential, raising the possibility that metabolic stress may become pathologically embedded through stable chromatin remodeling^37^. Chromatin-modifying enzymes are tightly coupled to cellular metabolic state^38,39^, and disruptions in core metabolic pathways may drive persistent epigenetic reprogramming that reinforces disease - associated cellular stress.

Here, we investigate the role of ATP13A2 LOF in astrocytes using human pluripotent stem cell-derived midbrain astrocytes. We demonstrate that ATP13A2 deficiency leads to lysosomal polyamine sequestration, impairing lysosomal function and triggering compensatory upregulation of *de novo* PA biosynthesis. Because astrocytes serve as a major cellular reservoir for polyamines and regulate their storage, disruption of lysosomal polyamine trafficking might be particularly deleterious. The metabolic rewiring depletes S-adenosyl methionine (SAM), a key methyl donor for DNA and histone methylation, resulting in widespread epigenetic remodeling and reprogramming of astrocytes towards a neurotoxic, inflammatory state that releases neurotoxic factors, including CXCL1, which leads to selective loss of dopaminergic neurons.

We identify AMD1, a key enzyme in PA biosynthesis, as a potential therapeutic target. Inhibiting AMD1 restores SAM levels, normalizes astrocyte epigenetic and lysosomal function, and mitigates astrocyte-mediated neurotoxicity. Together, our findings uncover a metabolically driven epigenetic mechanism linking lysosomal dysfunction to astrocyte-mediated dopaminergic loss, highlighting PA metabolism as a modulator of pathological aging and a promising target for therapeutic intervention in PD.

## RESULTS

### ATP13A2 LOF in astrocytes is neurotoxic to dopaminergic neurons

To investigate the impact of ATP13A2 loss-of-function (LOF) on dopaminergic neuron survival, we used isogenic human iPSC lines from a healthy male donor (ATP13A2^WT^)^40^ and a CRISPR-edited line harboring the ATP13A2 c.1306 loss-of-function mutation (ATP13A2^c1306^). The c.1306 G>A mutation introduces a frameshift resulting in a functional knockout of ATP13A2 and is causally associated with Kufor-Rakeb syndrome, an autosomal recessive form of juvenile Parkinsonism^41^ (**Fig. S1a-b**). Both ATP13A2^WT^ and ATP13A2^c1306^ iPSCs were differentiated into midbrain organoids containing tyrosine hydroxylase (TH)-positive dopaminergic neurons that recapitulate key molecular and cellular features of the *substantia nigra*^33,42^. Importantly, expression of canonical midbrain specification markers (*FOXA2*, *LMX1A*, and *NURR1*) was comparable between genotypes, indicating intact early midbrain patterning and enabling downstream analyses of disease-relevant phenotypes independent of developmental specification defects (**Fig S1c**).

In midbrain organoids, astrocytes emerge at approximately day 60 of differentiation and increase in abundance thereafter^33,42–44^. Consistent with intact early neuronal development, dopaminergic neuron marker TH was present at similar levels in ATP13A2^WT^ and ATP13A2^c1306^ organoids by day 25, before astrocyte appearance (p = 0.5513) (**Fig S1d**). In contrast, by day 100, after the appearance of astrocytes, ATP13A2^c1306^ organoids exhibited a significant reduction of immunoreactivity for dopaminergic neurons markers including TH and GIRK2 compared to isogenic ATP13A2^WT^ midbrain organoids (TH: Clone1: p = 0.0038, fold change 0.264±0.397; Clone2: p =0.0004, fold change 0.132±0.322) (GIRK2: p = 0.0149; fold change 0.652±0.373) (**Fig 1a and S1e**). At this stage, astrocytes populate the organoids, and we detected increased GFAP immunoreactivity in one ATP13A2 ^c1306^ clone compared to isogenic controls (Clone1: p = 0.0007, fold change 2.023±0.788) (**Fig 1b**). Together, these temporally resolved findings indicated that dopaminergic neuron loss in ATP13A2 LOF organoids emerges after maturation and coincides with astrocyte expansion, consistent with a potential non-cell-autonomous contribution to neurodegeneration.

**Figure 1.**
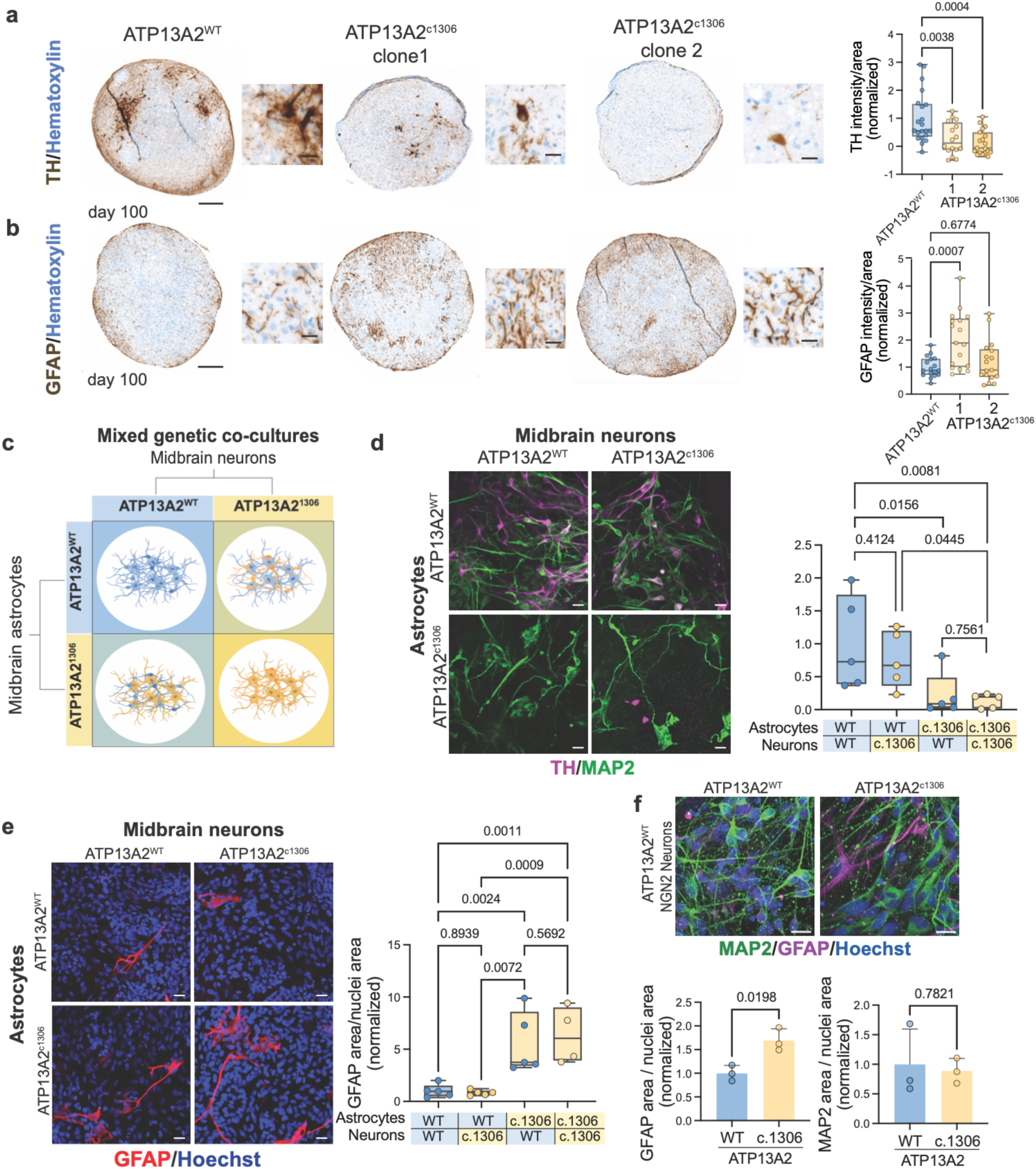
a. Micrography of TH (brown) and Hematoxylin (blue) staining for ATP13A2^WT^ and two clones of ATP13A2^c.1306^ day 100 midbrain organoids and quantification. Scale bar 200µm. Zoom-in scale 20µm; differences analyzed by One-way ANOVA. **b.** Micrography GFAP (brown) and Hematoxylin (blue) staining for WT and two clones of ATP13A2^c.1306^ day 100 midbrain organoids and quantification. Scale bar 200µm Zoom-in scale 20µm; differences analyzed by One-way ANOVA. **c.** Schematic representation of mixed genetic co-cultures of midbrain neurons and midbrain astrocytes. **d.** Immunocytochemistry of mixed genetic co-cultures of midbrain neurons and midbrain astrocytes stained for TH (magenta), MAP2 (green) and their quantification. Scale 50µm; differences analyzed by Two-way ANOVA. **e.** Immunocytochemistry of mixed genetic co-cultures of midbrain neurons and midbrain astrocytes stained for GFAP (red), and Hoechst (blue), and their quantification. Scale 50µm; differences analyzed by Two-way ANOVA. **f.** Immunocytochemistry of co-cultures of NGN2-cortical neurons and midbrain astrocytes stained for MAP2 (green), GFAP (magenta), and Hoechst (blue) and their quantification. Scale 20µm; differences analyzed by t-test.

To further investigate the role of astrocytes, we isolated midbrain astrocytes from day 100 ATP13A2^WT^ and ATP13A2^c1306^ organoids using previously described methods^33^. Isolated astrocytes exhibited comparable morphologies and homogenous expression of midbrain and astrocytic markers (S100b, SOX9, and FOXA2) (**Fig S1f-g)**. We then performed a combinatorial genetic mixing experiment by co-culturing ATP13A2^WT^ and ATP13A2^c1306^ midbrain neurons with either ATP13A2^WT^ or ATP13A2^c1306^ astrocytes (**Fig 1c**). Consistent with the organoid phenotypes, co-cultures composed entirely of ATP13A2^c1306^ cells showed a significant reduction in TH-positive neurons compared to all-ATP13A2^WT^ co-cultures (p = 0.0081, fold change 0.101±0.210) (**Fig 1d**). Strikingly, ATP13A2^WT^ neurons co-cultured with ATP13A2^c1306^ astrocytes phenocopied this loss of TH immunoreactivity, suggesting that ATP13A2-deficient astrocytes are sufficient to cause dopaminergic neuron degeneration (p = 0.0156, fold change 0.224± 0.408). In contrast, ATP13A2^c1306^ neurons co-cultured with ATP13A2^WT^ astrocytes retained a similar number of TH-positive neurons that was comparable to all-ATP13A2^WT^ co-cultures, indicating that loss of ATP13A2 in neurons alone is not sufficient and that astrocytic ATP13A2 loss is necessary for the neurodegeneration phenotype (p = 0.4124) (**Fig 1d**). Across co-culture conditions, the total astrocyte abundance, quantified as CD44-normalized area, did not differ significantly, suggesting that astrocyte abundance was not a confounding variable (vs. all-ATP13A2^WT^: ATP13A2^c1306^ neurons p = 0.9908; ATP13A2^c1306^ astrocytes p = 0.3627; all-ATP13A2^c1306^ p = 0.2367) (**Fig S1h**). However, notably, co-cultures containing ATP13A2^c1306^ astrocytes exhibited a significant increase in GFAP, even when co-cultured with WT-neurons, indicating that ATP13A2 loss promotes an astrocyte-autonomous reactive state (p = 0.0024; fold change 5.549±3.143) (**Fig 1e**).

Dopaminergic neurons comprise a minority of the neurons generated in midbrain cultures (∼5-20%^42,45^), raising the possibility that the observed phenotype reflects selective, rather than global, neuronal loss. To assess neuronal subtype specificity, we quantified total neuronal abundance using MAP2 immunoreactivity and evaluated neuronal survival. Across all midbrain co-culture conditions, the number of MAP2^+^ immunoreactive cells did not differ significantly, suggesting that ATP13A2^c1306^ astrocytes may be selectively toxic to dopaminergic neurons rather than inducing a generalized neuronal toxicity (vs. all-ATP13A2^WT^: ATP13A2^c1306^ neurons p = 0.8136; ATP13A2^c1306^ astrocytes p = 0.7970; all-ATP13A2^c1306^ p = 0.1136) (**Fig S1h**). To specifically test this, we co-culture ATP13A2 isogenic astrocytes with NGN2-induced cortical neurons that lack dopaminergic neuron subtypes. In this context, ATP13A2^c1306^ astrocytes again exhibited significantly increased GFAP immunoreactivity, consistent with a cell-autonomous reactive phenotype (staining p = 0.0198, fold change 1.694±0.647; immunoblot p = 0.0164, fold change 5.245±2.745) (**Fig 1f, Fig S1i**). Notably, this reactive state did not impair cortical neuron survival, as indicated by no significant change in the number of MAP2 -positive neurons, and protein levels for the pan-neuronal marker β-neuronal tubulin (TUJ1) were modestly increased compared to isogenic controls (MAP2: p = 0.7821; TUJ1: p = 0.0347, fold change 1.444±0.450) (**Fig 1f and S1i**). Together, these data demonstrate that ATP13A2 LOF confers a cell-autonomous state marked by reactive astrocyte signatures that is selectively toxic to dopaminergic neurons, while sparing other neuronal populations.

### ATP13A2 LOF in astrocytes alters lysosomal homeostasis and promotes inflammation

We observed that GFAP is consistently elevated in ATP13A2^c1306^ astrocytes in co-cultures, suggesting that ATP13A2 ^c1306^ astrocytes have reactive inflammatory properties. To isolate cell-autonomous effects, we next assessed markers of astrocyte inflammation in monocultures (**Fig 1b,e,f**). Immunostaining revealed significant upregulation of multiple reactive astrocyte markers that overlap with signatures of astrocyte reactivity observed in the *substantia nigra* of idiopathic PD patients^46^, including GFAP, CD44, and S100A6, in ATP13A2^c1306^ astrocytes compared to isogenic ATP13A2^WT^ astrocytes (GFAP: p = 0.0002, fold change 3.628±2.196; CD44: p = 0.0015, fold change 1.433±0.597; S100A6: p = 0.0258, fold change 1.202±0.313) (**Fig 2a**). To systematically investigate the molecular program underlying this inflammatory state, we performed bulk RNA sequencing on isogenic ATP13A2^WT^ and ATP13A2^c1306^ astrocytes. Differential expression analysis identified 228 upregulated and 201 downregulated genes in the ATP13A2^c1306^ astrocytes compared to isogenic control astrocytes (absolute Log_2_FC>0.7; FDR<0.01). Gene ontology and pathway enrichment analysis of the upregulated genes showed significant enrichment of pathways associated with cytokine signaling and extracellular matrix remodeling, consistent with a pro-inflammatory reactive astrocyte phenotype (**Fig S2a**)^47,48^. Notably, ATP13A2^c1306^ astrocytes showed robust induction of inflammatory cytokines (*IL11, IL16, IL32, IL34*), chemokines (*CXCL1, CXCL14, CXCL6, CXCL8, TAFA5*), and disease-associated astrocyte markers such as *AMIGO2* and *GFAP* (**Fig 2b**, orange labels). In contrast, genes associated with lysosomal function, including multiple lysosomal proteases (*CTSB, CTSZ*), were significantly downregulated in ATP13A2^c1306^ compared to isogenic control astrocytes, suggesting impaired lysosomal clearance in ATP13A2-deficient astrocytes (**Fig 2b**, green labels). Together, these data support a model in which ATP13A2 LOF disrupts lysosomal function in astrocytes, triggering a transcriptional shift towards a pro-inflammatory reactive state. To directly test whether impaired lysosomal polyamine export is sufficient to induce astrocyte reactivity, we pharmacologically blocked lysosomal polyamine export in ATP13A2^WT^ astrocytes using AMXT-1501, a PA transporter inhibitor^49,50^. This treatment led to significant upregulation of key transcriptional signatures found in ATP13A2^c1306^ astrocytes, inducing *GFAP, S100A6, AMIGO2*, and *CXCL1*, thereby establishing polyamine lysosomal retention of polyamines as a causal upstream driver of astrocyte reactivity (**Fig S2b**).

**Figure 2.**
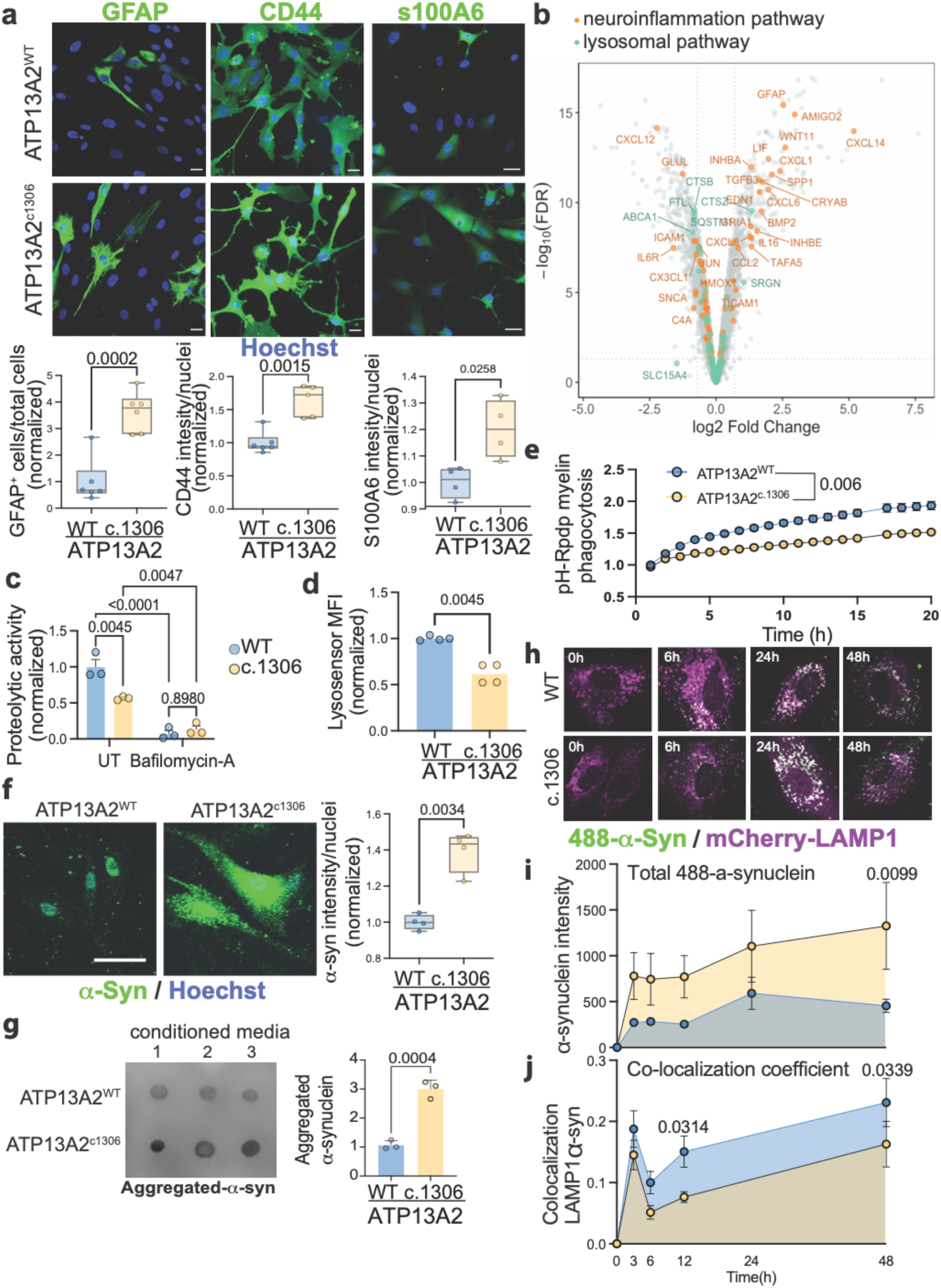
a. Immunocytochemistry of astrocytes labeled for GFAP, CD44, and S100A6 and quantification. Scale bar 20µm. Differences were analyzed by a two-tailed t-test. **b.** Volcano plot of RNAseq results, evidenced with colors genes in neuroinflammatory pathways (orange) and lysosomal pathways (green), only genes with an absolute log2FC > 0.7 are labeled. **c.** quantification of the area under the curve of DQ-BSA lysosome proteolysis assay signal over 20h of acquisition. Bafilomicyn-A is used as a control to block lysosomal function. Differences were analyzed by Two-way ANOVA. **d.** Median fluorescence intensity of Lysosensor signal in ATP13A2^WT^ and ATP13A2^c.1306^ astrocytes. Differences were analyzed by a two-tailed t-test. **e.** Quantification of Red Fluorescent intensity signal over 20h of acquisition of pH-rodo-myelin signal, normalized by cell confluence. Differences analyzed by Two-tailed t-test. **f.** Immunocytochemistry of astrocytes labeled for α-synuclein and quantification. Scale bar 20µm. Differences were analyzed by a two-tailed t-test. **g.** Dot blot of astrocyte-conditioned media blotted with aggregated α-synuclein antibody and quantification. Differences were analyzed by a two-tailed t-test. **h.** Live imaging time course representative images of mCherry-LAMP1 signal (magenta) and recombinant α-synuclein-HiLyte (488-synuclein, green). **i.** Quantification of total 488- α-synuclein intensity normalized by cell count and **j.** Mander’s co-localization coefficient of green colocalizing with magenta signal over time. Differences were analyzed by Two-way ANOVA.

Impaired lysosomal function is a central contributor to α-synuclein accumulation and dopaminergic neuron loss in PD^3,33,51–53^. Consistent with this, lysosomal proteolysis activity, measured by DQ-BSA cleavage, was reduced by ∼40% in ATP13A2^c1306^ astrocytes compared to isogenic controls, indicating a marked deficit in lysosomal degradation capacity (p = 0.0045) (**Fig 2c**). Complementing this finding, LysoSensor measurements showed significantly reduced signal in ATP13A2^c1306^ astrocytes, indicating de-acidification (elevated intra-lysosomal pH), relative to ATP13A2^WT^ controls (p = 0.0045, fold change 0.6182±0.243) (**Fig 2d**). Functional consequences of this lysosomal impairment were further evident in reduced phagolysosome delivery, assessed using pH-rodo-labeled myelin debris, which depends on trafficking to and fusion with acidic lysosomes (**Fig 2e**). Pharmacologic inhibition of lysosomal polyamine export using AMXT-1501 in ATP13A2^WT^ astrocytes also reduced both DQ-BSA proteolysis and LysoSensor signal (**Fig S2c-d**), establishing that impaired ATP13A2-dependent polyamine transport disrupts lysosomal acidification and impairs proteolytic function in astrocytes. Notably, despite impaired lysosomal function, ATP13A2^c1306^ astrocytes, and AMXT-1501-treated ATP13A2^WT^ both exhibited a significant increase in lysosomal mass, as assessed by LAMP1 immunostaining (vs. ATP13A2^WT^ UT, ATP13A2^c1306^ p = 0.114, fold change 1.482±0.093; ATP13A2^WT^ AMXT-1501 p = 0.114, fold change 1.492±0.146) (**Fig S2e**). This phenotype is consistent with previous reports in ATP13A2-deficient cellular and animal models^4,27,54,55^, indicating that impaired polyamine transport triggers lysosomal accumulation, but fails to support proper lysosomal acidification and proteolytic function, resulting in an expanded, yet dysfunctional lysosomal compartment in astrocytes.

Because lysosomes are the primary site for α-synuclein degradation, we next examined α-synuclein handling in ATP13A2^c1306^ astrocytes^56–60^. Immunofluorescence and immunoblot analyses revealed a significant increase in α-synuclein protein in ATP13A2^c1306^ astrocytes compared to isogenic controls (p = 0.0034, fold change 1.393±0.313) (**Fig 2f, S2f**). However, increased α-synuclein occurred without changes in *SNCA* transcripts abundance, as assessed by RNA-seq and qPCR, suggesting that increased α-synuclein in astrocytes was a result of impaired degradation rather than transcriptional upregulation (**Fig S2g**). Consistent with defective clearance, conditioned media from ATP13A2^c1306^ astrocytes contained significantly higher levels of aggregated α-synuclein than isogenic controls (p = 0.0004, fold change 2.884±1.084) (**Fig 2g**). To directly assess intracellular trafficking to the lysosomal compartment, we performed live-cell imaging of mCherry-LAMP1-labeled lysosomes^61^ following exposure to fluorescently tagged α-synuclein monomers (**Fig S2h)**. Compared to isogenic control astrocytes, ATP13A2^c1306^ astrocytes exhibited a significant increase in total intracellular α-synuclein accumulation at all timepoints considered, accompanied by a marked reduction in α-synuclein localization within LAMP1-positive signal (**Fig 2h-j**). These findings indicate that ATP13A2 LOF impairs α-synuclein targeting and delivery to the lysosomal compartment, consistent with defective lysosome-mediated clearance.

### The Secretome of ATP13A2 LOF astrocytes is altered and contains factors that cause dopaminergic neurotoxicity

Our findings suggest that ATP13A2 LOF astrocytes may drive neurodegeneration through at least two non-mutually exclusive mechanisms: impaired intracellular clearance of pathogenic proteins and debris, and secretion of neurotoxic factors. To isolate the specific contribution of astrocyte-secreted signals, we performed astrocyte-conditioned media (ACM) experiments (**Fig 3a**).

**Figure 3.**
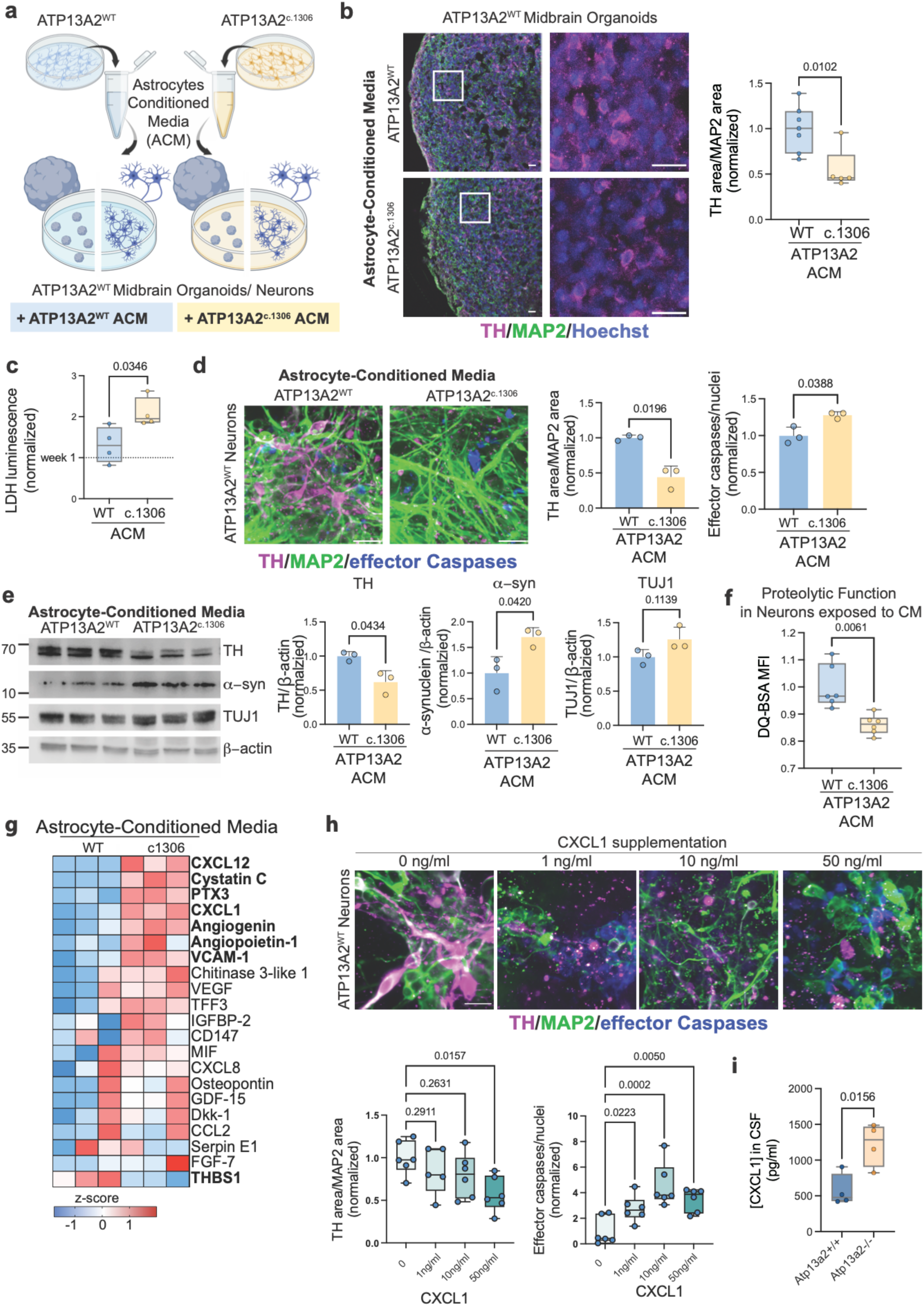
a. Schematic representation of astrocyte conditioned media (ACM) experiments: media exposed to 2D astrocyte cultures were collected every 4 days and used to culture ATP13A2^WT^ midbrain organoids or ATP13A2^WT^ midbrain neuronal cultures. **b.** ATP13A2^WT^ midbrain organoids exposed to ACM and quantification. Staining for TH (magenta), MAP2 (green), and Hoechst (blue). Scale bar 50µm. Differences were analyzed by a two-tailed t-test. **c.** LDH assay ACM experiments on ATP13A2^WT^ midbrain neurons at week four. Values are normalized to ATP13A2^WT^ week one levels (horizontal line); differences analyzed by Two-way ANOVA. **d.** Immunofluorescence of TH (magenta), MAP2 (green), and active (cleaved) effector caspase 3/7 (blue) in ATP13A2^WT^ midbrain neurons exposed to ACM. Differences were analyzed by a two-tailed t-test. **e.** Immunoblots of ATP13A2^WT^ midbrain neurons exposed to conditioned media and quantification of dopaminergic neurons (TH), α-synuclein (α-syn), and total neurons (TUJ1). ®-Actin was used as a loading control. Differences were analyzed by a two-tailed t-test. **f.** Median fluorescence intensity of DQ-BSA to measure proteolytic function of ATP13A2^WT^ neurons exposed to conditioned media. Differences were analyzed by a two-tailed t-test. **g.** Heat map showing the relative levels of cytokines detected in astrocyte-conditioned media. Cytokines in bold are statistically different by Two-way ANOVA (p < 0.05). **h.** Immunofluorescence of TH (magenta), MAP2 (green), and active (cleaved) effector caspase 3/7 (blue) in neurons exposed to WT conditioned media supplemented with increasing amount CXCL1. Differences were analyzed by One-way ANOVA.

Initially, day 25 ATP13A2^WT^ 3D midbrain organoids were exposed to conditioned media collected from either ATP13A2^WT^ or ATP13A2^c1306^ astrocytes. Treatment with ATP13A2^c1306^ ACM resulted in a reduction in the number of tyrosine hydroxylase (TH)-positive dopaminergic neurons (p = 0.0102, fold change 0.548±0.363) (**Fig 3b**). To further characterize this neurotoxic effect, we exposed day 25 ATP13A2^WT^ 2D midbrain neuronal cultures to the same ACM variants. We observed a time-dependent increase in cell death; while no significant changes were detected at earlier time points (**Fig S3a**), four weeks of exposure to ATP13A2^c1306^ ACM resulted in a significant elevation of extracellular lactate dehydrogenase (LDH), indicating a loss of membrane integrity (p = 0.0346, fold change 1.596±0.916) (**Fig 3c**). This was accompanied by a marked increase in cleaved effector caspase activity, which we measured through Image-IT effector cleaved caspase staining, confirming that the observed cell death was apoptotic (p = 0.0388, fold change 1.278±0.368) (**Fig 3d**). Consistent with our 3D organoid data, this neurotoxicity was selective for dopaminergic neurons, as evidenced by decreased TH/MAP2 immunostaining in the 2D cultures (p = 0.0196, fold change 0.443±0.306) (**Fig 3d**). Furthermore, neurons exposed to ATP13A2^c1306^ ACM exhibited an accumulation of α-synuclein and a reduction in DQ-BSA-derived fluorescence compared to ATP13A2^WT^ ACM, indicating that astrocyte-derived factors induce a secondary impairment of neuronal lysosomal proteolytic capacity. (α-synuclein p = 0.0420, fold change 1.706±0.802; DQ-BSA MFI p = 0.0061, fold change 0.861±0.198) (**Fig 3e-f**).

To identify candidate astrocyte-derived mediators of neurotoxicity, we profiled ACM using an unbiased cytokine array. Consistent with transcriptional changes, ATP13A2^c1306^ ACM was significantly enriched for inflammatory factors (CXCL8, CXCL1, CCL2, MIF, PTX3), angiogenic factors (VEGF, Angiogenin, Angiopoietin-1, FGF-7), and extracellular matrix and tissue remodeling proteins (CHI3L1, THBS1, Osteopontin), highlighting a complex, disease-relevant secretome (**Fig 3g**). Among these factors, CXCL1 was consistently elevated across replicates. We validated this increase quantitatively via ELISA, confirming that CXCL1 protein levels are significantly higher in ATP13A2^c1306^ ACM compared to control ACM (**Fig S3b**). Given that the CXCL1-CXCR2 signaling axis has been previously implicated in neurodegeneration and systemic CXCL1 is sufficient to induce dopaminergic neuron loss in the substantia nigra *in vivo*^62,63^, we therefore hypothesized that CXCL1 could be a primary driver of the observed phenotype.

To test this, we supplemented ATP13A2^WT^ midbrain neuronal cultures with recombinant CXCL1 at increasing concentrations (0, 1, 10, 50, and 100 ng/ml). We observed a clear dose-dependent trend in apoptotic activation indicated by a significant increase in cleaved caspase-3 immunoreactivity at 50 and 100 ng/ml of CXCL1 (**Fig 3h, S3c**). Notably, CXCL1 toxicity exhibited a window of selectivity for dopaminergic neurons. 50 ng/ml of CXCL1 significantly reduced the abundance of dopaminergic (TH-positive) neurons without decreasing total neuronal content (MAP2/nuclei), recapitulating the preferential vulnerability seen in our original ACM experiments (TH/MAP2 p = 0.0157, fold change 0.681±0.166) (**Fig 3h**). In contrast, the 100 ng/ml dose resulted in broad, non-selective neurotoxicity (**Fig S3d**). To assess whether CXCL1 is a soluble factor associated with ATP13A2 LOF in vivo, we employed Atp13a2 knock-out (-/-) mice. Consistent with these *in vitro* findings, the concentration of Cxcl11 in cerebrospinal fluid (CSF) was 2-fold higher in Atp13a2-/- knock-out mice compared to control animals (Atp13a2+/+) (p = 0.0156, fold change 2.137±0.501) (**Fig 3i**). Together, these data demonstrate that CXCL1 is both sufficient to induce selective dopaminergic neuron loss *in vitro* and elevated *in vivo* in the context of ATP13A2 deficiency, supporting a role for CXCL1 as a relevant secreted mediator of dopaminergic neuron toxicity.

### ATP13A2 LOF alters the bioavailability of SAM, leading to epigenetic reprogramming of astrocytes to a neuroinflammatory state

Polyamines enter cells primarily through endolysosomal trafficking, where they transiently accumulate in the lysosome. ATP13A2 is a critical lysosomal polyamine exporter with a high affinity for spermine (SPM) and spermidine (SPD), thereby maintaining proper cytosolic polyamine distribution^5^. We measured cytosolic polyamine levels using a doxycycline-inducible, genetically encoded reporter^64^. This reporter utilizes a polyamine-responsive +1 frameshifting element where mCherry serves as a normalization control, and eYFP is expressed only upon a polyamine-dependent frameshift. In ATP13A2^c1306^ astrocytes, we observed a significant reduction in the eYFP/mCherry fluorescence ratio compared to isogenic controls, indicating depleted polyamine levels (p = 0.0001, fold change 0.738±0.224) (**Fig 4b**). Complementary uptake assays using labeled BODIPY-SPM and BODIPY-SPD showed a modest but significant increase in polyamine influx in ATP13A2^c1306^ astrocytes (SPM: p = 0.0276; SPD: p = 0.0276) (**Fig S4a**). Consistent with previously reported findings^5^, we observed that the cytosol depletion of polyamines in ATP13A2^c.1306^ astrocytes results from their increased sequestration within the lysosomal compartment. Specifically, both BODIPY-SPM and BODIPY-SPD showed significantly increased co-localization with LAMP1-positive lysosomes compared to control astrocytes (SPM: p = 0.0085; SPD: p = 0.0085) (**Fig 4c**).

**Figure 4.**
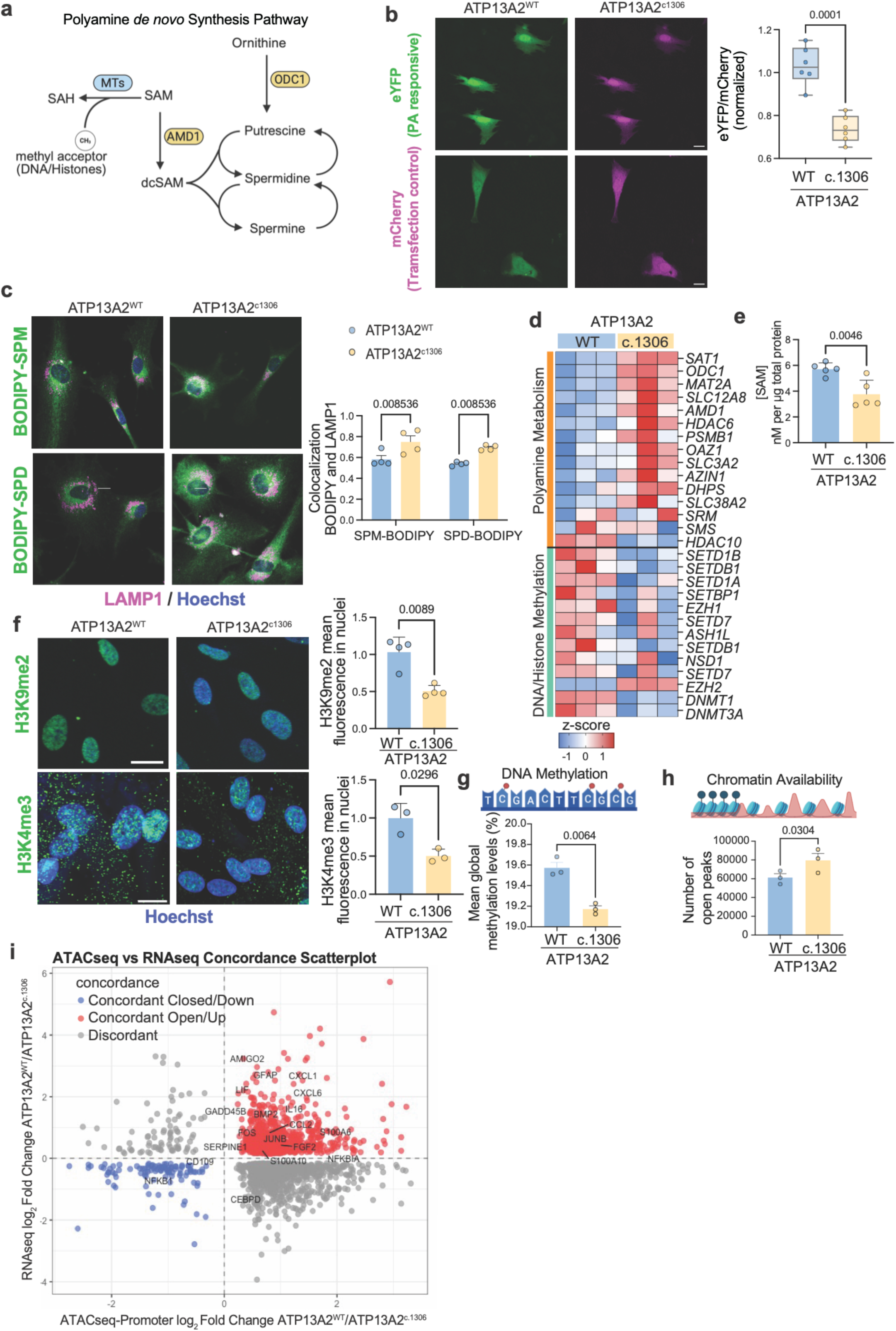
a. Schematic representation of the PA *de novo* synthesis pathway. **b.** Astrocytes transduced with PA sensors, and quantification of YFP/mCherry signal. Scale bar 20µm. Differences were analyzed by Two-tailed t-test. **c.** Astrocytes exposed to BODIPY-polyamine (green), stained for LAMP1 (magenta), nuclei (blue) and co-localization coefficient quantification. Differences were analyzed by Two-way ANOVA. **d.** heatmap of genes detected by bulk RNA-seq involved in PA synthesis (orange label) and DNA/Histone methylation (green label). **e.** SAM quantification by ELISA. Differences analyzed by Two-tailed t-test. **f.** Immunofluorescence of histone-3 methylation marks (green) and Hoechst (blue): K9me2 for inactive chromatin, K4me3 for active chromatin. Differences were analyzed by a two-tailed t-test. **g.** Schematic of DNA methylation and global methylation levels detected by whole genome bisulphate sequencing. Differences were analyzed by a two-tailed t-test. **h.** schematic of ATAC-seq peaks and quantification of the number of peaks. Differences were analyzed by a two-tailed t-test. **h.** Concordance plot between ATACseq and RNAseq analysis, neuroinflammatory genes are labeled in the plot.

In the cytosol, polyamines are essential for cellular signaling and mitochondrial function^13,14^. Therefore, we hypothesize that increased lysosomal retention and the resulting reduction of cytosolic polyamines in ATP13A2^c.1306^ astrocytes could lead to compensatory upregulation of *de novo* polyamine biosynthesis. Consistent with this, RNA-seq analysis of the *de novo* polyamine biosynthetic pathway (schematized in **Fig 4a**) revealed coordinated induction of polyamine synthesis and transport genes in ATP13A2 LOF astrocytes, including *ODC1, AMD1, SAT1, ATP13A3,* along with *MAT2A,* which encodes the SAM-synthesizing enzyme required to sustain elevated polyamine demand (**Fig 4d**, orange label). qPCR validation confirmed upregulation of key pathway nodes (*ODC1* 1.521±0.104; *AMD1* 1.921±0.252) (**Fig s4b**). The *de novo* synthesis of SPM and SPD requires two molecules of S-Adenosyl-methionine (SAM)^65^ (**Fig 4a**). Consistent with increased flux through the SAM-dependent polyamine synthesis pathway, quantification of SAM by ELISA showed a significant reduction in intracellular SAM in ATP13A2^c1306^ astrocytes relative to isogenic controls (p = 0.0046, fold change 0.66±0.088) (**Fig 4e**).

SAM is the principal methyl-donor for DNA and histone methylation^66–68^. Therefore, the use of SAM for polyamine synthesis could deplete its bioavailability for DNA and histone methylation. In line with constrained methyl-donor availability, we observed a broad and significant reduction in the expression of genes associated with DNA and histone methylation in ATP13A2^c1306^ astrocytes (**Fig 4d**, green label). The downregulation included key DNA methyltransferases (*DNMT1, DNMT3A*) and histone methyltransferases, notably members of the SET domain-containing family (*SEDTD1A, SETD1B, SETDB1, SETD7*) and the polycomb repressive complex component *EZH1*. Additionally, other epigenetic regulators such as NSD1*, ASH1L,* and *SETBP1,* were also reduced. Collectively, these transcriptomic changes suggest that polyamines and subsequent consumption of SAM lead to a systemic disruption of the cellular methylation landscape.

Consistent with impaired methylation capacity, global levels of both H3K9me2, a repressive heterochromatin-associated mark, and H3K4me3, an active promoter-associated mark, were significantly decreased in ATP13A2^c1306^ astrocytes relative to isogenic controls (H3K9me2: p = 0.0296, fold change 0.743±0.200; H3K4me3: p = 0.0089, fold change 0.504±0.281) (**Fig 4f)**. Similarly, whole genome bisulfite sequencing (WGBS) corroborated this trend, revealing a widespread CpG hypomethylation across the genome of ATP13A2^c1306^ astrocytes compared to isogenic control astrocytes (**Fig 4g).** We focused exclusively on proximal promoter regions (≤2kb from the transcription starting site), given the well-established relationship between promoter DNA methylation and transcriptional activity^69,70^. Using stringent thresholds (absolute methylation difference>0.3, q value<0.01), we identified 1,662 hypomethylated and 1,406 hypermethylated promoters in ATP13A2^c.1306^ astrocytes relative to isogenic controls. Volcano plot highlighted a few neuroinflammatory genes among the hypomethylated promoters, including *CXCL1* (**Fig S4c**).

To assess the functional consequences of these epigenetic changes, we performed ATAC-seq profiling, which identified extensive chromatin remodeling in ATP13A2^c.1306^ astrocytes. More than 18,000 genomic regions exhibited increased chromatin accessibility compared to controls (**Fig 4h**). Although ATP13A2^c1306^ astrocytes showed a small reduction in overall promoter accessibility genome-wide (39.6% vs 43.25% in ATP13A2^WT^, **Fig S4d**), regions that became differentially accessible in the mutant condition were disproportionately enriched at promoters (approximately 40% vs. 20% in ATP13A2^WT^), indicating preferential promoter opening rather than uniform chromatin relaxation (**Fig S4e)**.

Integration of the ATAC-seq and RNA-seq datasets revealed concordance between chromatin accessibility and transcriptional activation. Genes upregulated in ATP13A2^c.1306^ astrocytes were significantly enriched among loci with increased chromatin accessibility, particularly at neuroinflammatory and astrocyte reactivity genes (labeled in **Fig 4h**). Locus-level inspection further validated these findings, revealing increased accessibility specifically at core inflammatory chemokines (*CXCL1, CXCL2, CXCL3, CXCL6, CXCL12, CCL2*) and astrocyte reactivity genes (*GFAP, AMIGO2, VIM, SERPINA3*) (**Fig S4f**).

Together, these data demonstrate that ATP13A2 loss in astrocytes sequesters polyamines within lysosomes, thereby depleting cytosolic polyamines and inducing compensatory upregulation of polyamine biosynthesis, which reduces cytosolic SAM levels. This metabolic rewiring induces widespread DNA hypomethylation and chromatin opening and epigenetically primes astrocytes for a neuroinflammatory transcriptional state.

### Inhibiting AMD1 Restores SAM Levels, Lysosomal Function, and Attenuates Astrocyte-Mediated Neurotoxicity

Given that ATP13A2 LOF astrocytes exhibit SAM depletion and an inflammatory state associated with epigenetic changes, we asked whether reducing SAM consumption through polyamine biosynthesis could reverse these phenotypes. To test this, we inhibited AMD1, the enzyme responsible for converting SAM into decarboxylated SAM (dcSAM), using methylglyoxalbisguanylhydrazone (MGBG) in ATP13A2^c1306^ astrocytes. MGBG treatment significantly increased intracellular SAM levels (fold change 2.713±0.614, p = 0.0254) ( **Fig 5a**), indicating effective suppression of SAM decarboxylation. Restoration of SAM was accompanied by recovery of lysosomal proteolytic activity, as measured by DQ-BSA processing (ATP13A2^c1306^ MGBG vs control fold change 1.963±0.07, p < 0.0001) (**Fig 5b**). Critically, MGBG also reversed the reactive astrocyte phenotype, reducing GFAP expression at both the transcript and the protein levels (ATP13A2^c1306^ MGBG vs control fold change 0.23±0.144, p = 0.0281) (**Fig 5c, S5a**). Consistent with this, MGBG treatment significantly decreased the concentration of pro-inflammatory chemokine CXCL1 in ATP13A2^c.1306^ astrocyte-conditioned media (fold change 0.434±0.311, p = 0.0179) (**Fig 5d**). AMD1 inhibition restored the repressive chromatin landscape mark H3K9me2, increasing the levels that were reduced in ATP13A2^c1306^ astrocytes (ATP13A2^c1306^ MGBG vs control fold change 1.261±0.478, p = 0.0464) (**Fig 5e**).

**Figure 5.**
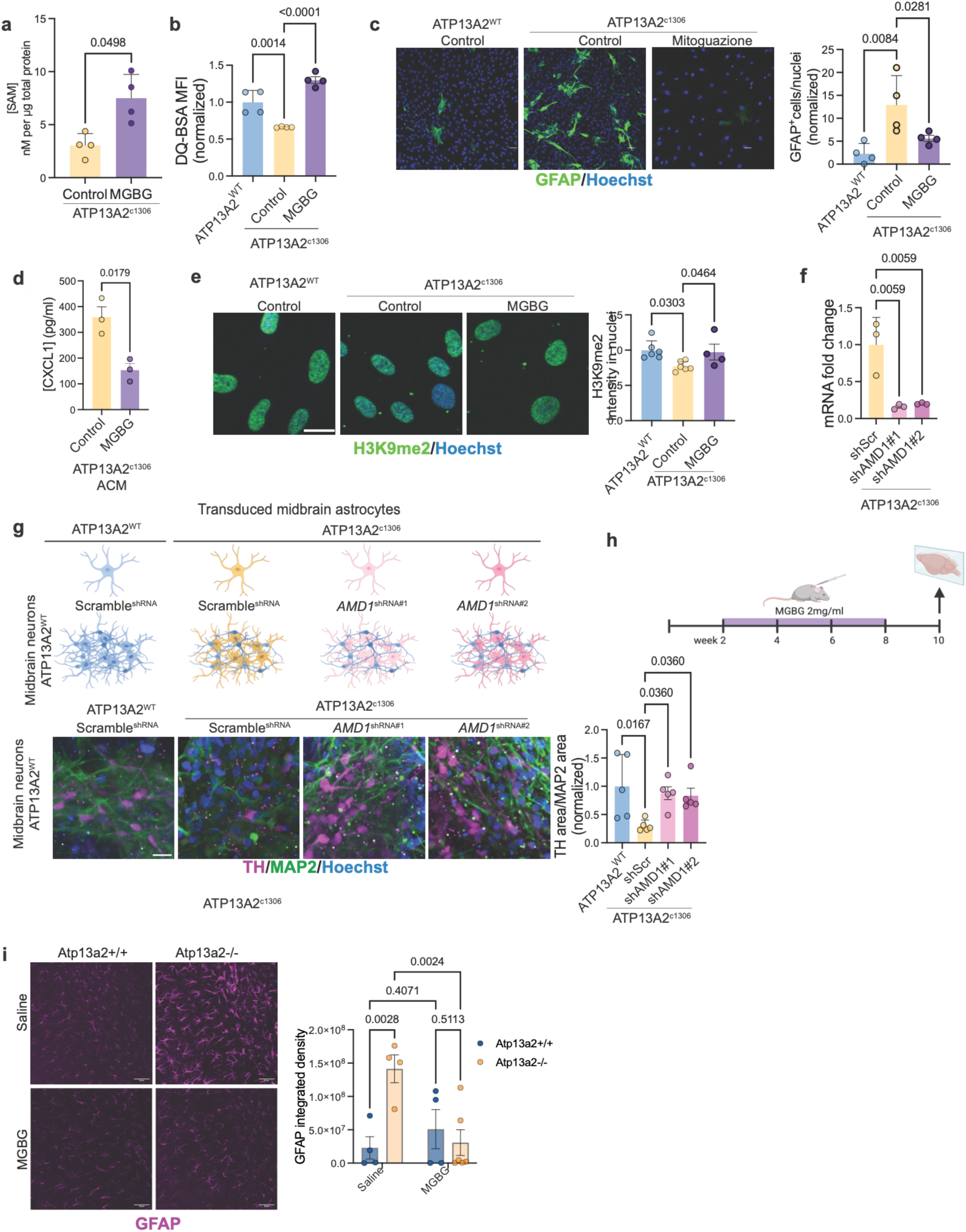
a. SAM quantification by ELISA. Differences were analyzed by a two-tailed t-test. **b.** Median fluorescence intensity of DQ-BSA in ATP13A2^WT^ and ATP13A2^c.1306^ astrocytes treated with MGBG or control solution (H2O). **c.** Immunocytochemistry of ATP13A2^WT^ and ATP13A2^c.1306^ astrocytes treated with MGBG or control solution (H2O), labeled for GFAP (green) and Hoechst (blue), and quantification. Differences were analyzed by One-way ANOVA. **d.** CXCL1 ELISA on conditioned media from astrocytes treated with MGBG and control. Differences were analyzed by a two-tailed t-test. **e.** Immunofluorescence of histone k9 dimethylation mark (green) and Hoechst (blue) in ATP13A2^WT^ and ATP13A2^c.1306^ astrocytes treated with MGBG or control solution (H2O). Differences were analyzed by One-way ANOVA. **f.** mRNA fold change by qPCR of *AMD1* in astrocytes targeted with shRNAs. **g.** Schematic representation of the co-culture experiment and immunocytochemistry of TH (magenta), MAP2 (green), and nuclei (blue), of co-cultures of midbrain neurons and astrocytes transduced with shRNA targeting AMD1 or control (shScramble), and quantification of dopaminergic neurons. Differences were analyzed by One-way ANOVA. **h.** schematic representation of treatments in mice **i.** Images of Atp13a2 knock-out (-/-) mice and control (+/+) brain slices stained for GFAP (magenta) and quantification of GFAP area and integrated intensity. Differences were analyzed by Two-way ANOVA.

To exclude off-target pharmacological effects, we next performed shRNA-mediated knockdown of *AMD1* in ATP13A2^c1306^ astrocytes. We identified two AMD1-shRNAs that achieved an approximately 80% reduction in *AMD1* expression compared to the scramble control shRNA (p = 0.0059) (**Fig 5f**). Similar to MGBG, shRNA knockdown of *AMD1* significantly increased lysosomal proteolytic activity in ATP13A2^c.1306^ astrocytes, achieving similar levels as seen in ATP13A2^WT^ astrocytes (vs ATP13A2^c1306^ shScr, shAMD1#1 fold change 1.317±0.206, p = 0.0432; shAMD1#2-fold change1.514±0.259, p = 0.0046) (**Fig S5b).** Likewise, shRNA knockdown of AMD1 significantly reduced reactive astrocyte marker GFAP compared to scramble control, again achieving similar levels of GFAP expression as WT control astrocytes (vs ATP13A2^c1306^ shScr, shAMD1#1 fold change 0.144±0.026, p < 0.0001; shAMD1#2-fold change 0.168±0.062, p < 0.0001) (**Fig S5c**).

We next assessed the functional consequences of astrocyte-specific AMD1 knockdown in midbrain neuron-astrocyte co-cultures. Astrocytes were transduced with the shAMD1 or shScramble as a control and then were co-cultured with ATP13A2^WT^ neurons. Consistent with our previous results, the co-culture with ATP13A2^c1306^ shScr resulted in a significant reduction in the number of TH-positive neurons compared to ATP13A2^WT^ shScr astrocytes (fold change 0.301±0.261, p = 0.0167). In contrast, ATP13A2^c1306^ astrocytes expressing independently two different shRNAs against AMD1 failed to induce dopaminergic neuron loss, preserving TH^+^ dopaminergic neuron abundance relative to shScr control (vs ATP13A2^c1306^ shScr, shAMD1#1-fold change 2.910±1.285, p = 0.036; shAMD1#2-fold change 2.774±1.263, p = 0.036) (**Fig 5g**). These results demonstrate that astrocyte-specific knockdown of AMD1 is sufficient to prevent dopaminergic neuron degeneration mediated by ATP13A2 loss-of-function astrocytes.

Finally, to evaluate the relevance of AMD1 inhibition *in vivo*, we took advantage of the Atp13a2 knockout mouse model, which has been reported to exhibit elevated astrocytic activity^27^. Consistent Atp13A2^-/-^ mice displayed robust GFAP immunoreactivity at baseline (Saline condition fold change 6.191±4.613, p=0.0028) (**Fig 5i**). Postnatal treatment with the AMD1 inhibitor MGBG (2mg/ml, P14-P56; **Fig 5h**) significantly reduced GFAP immunoreactivity in Atp13a2^-/-^ mice toward wild-type (Atp13a2^+/+^) levels (fold change 0.217±0.379, p=0.0024) (**Fig 5i**), indicating suppression of astrocyte reactivity *in vivo*. Collectively, our findings establish AMD1 as a central metabolic regulator of astrocyte homeostasis downstream of ATP13A2 loss and identify its inhibition as a strategy to restore SAM balance, reverse lysosomal dysfunction, and protect dopaminergic neurons.

## Discussion

Our findings establish a critical mechanism by which lysosomal polyamine dysregulation in astrocytes drives non-cell-autonomous dopaminergic neurodegeneration. Using isogenic human iPSC-derived midbrain models, we demonstrated that ATP13A2 LOF in astrocytes leads to lysosomal sequestration of polyamines, triggering compensatory *de novo* polyamine synthesis that depletes S-adenosylmethionine (SAM) and induces epigenetic remodeling. This metabolic-epigenetic rewiring stabilizes a reactive astrocyte state characterized by impaired lysosomal function and the secretion of neurotoxic inflammatory mediators. Among these, CXCL1 emerges as a functionally relevant component of the ATP13A2-deficient astrocyte secretome that is sufficient to induce selective dopaminergic neuron death.

Lysosomal dysfunction has been proposed as a central pathway in PD, with mutations in lysosome-associated genes defining a major axis of inherited risk and vulnerability^4,7,51,55^. However, how lysosomal defects are translated into stable, cell-type-specific pathogenic programs remains unclear. Our data suggest that, in astrocytes, lysosomal polyamine sequestration initiates a metabolic cascade that becomes epigenetically embedded through SAM depletion, thereby converting lysosomal stress into a persistent inflammatory state. We show that even acute pharmacological blockade of polyamine export with AMXT-1501 is sufficient to induce astrocytic neuroinflammatory responses, supporting a direct causal link between impaired polyamine trafficking and glial activation. Astrocytes might be particularly vulnerable to this mechanism, given their central role as major reservoirs and distributors of polyamines in the brain^34,71,72^, as well as their high metabolic and epigenetic plasticity^73–75^.

Chromatin-modifying enzymes are highly sensitive to metabolite availability, including SAM, and changes in cellular metabolism can directly shape chromatin landscapes^52,76–79^. The epigenetic consequences of SAM depletion observed in our model (global DNA hypomethylation, loss of repressive chromatin structure, and activation of inflammatory gene-expression programs) parallel molecular features of brain aging, which are established contributors to neurodegeneratio^68,80^. Similar metabolic-epigenetic coupling mechanisms may operate in other lysosomal and glial-driven neurodegenerative disorders, suggesting broader relevance beyond ATP13A2-associated PD.

Our results align and extend emerging frameworks that position astrocytes as active determinants in neuronal health, rather than passive responders to neuronal injury. Astrocytes have been shown to regulate neuronal function and viability through tightly controlled secretory programs^81–83^. Our data identify SAM and polyamine metabolism as upstream determinants of a neurotoxic secretory state in astrocytes. This provides a molecular explanation for the early gliosis observed in Atp13a2 animal models, in which astrocyte dysfunction precedes motor phenotypes^27^. Importantly, we extend these findings *in vivo* by showing that postnatal inhibition of AMD1 in *Atp13A2* knock-out mice suppresses astrocyte reactivity, reducing elevated GFAP immunoreactivity.

Conditioned media experiments revealed that astrocyte-induced neurotoxicity occurs through a non-cell autonomous mechanism mediated by secreted factors. Notably, both NGN2-induced cortical neurons and non-dopaminergic midbrain neurons in co-culture experiments were resistant to ATP13A2 LOF astrocytes’ conditioned media, highlighting the selective vulnerability of dopaminergic neurons. Moreover, our findings that ATP13A2^WT^ astrocytes restore dopaminergic neuronal survival when cocultured with ATP13A2 LOF neurons further support the critical role of astrocytes in shaping neuronal viability and reinforce the concept that astrocyte state, rather than neuronal genotype alone, is a key determinant for dopaminergic neuron survival. A key insight from our study is the identification of CXCL1 as a mechanistic effector of astrocyte-mediated toxicity. CXCL1-CXCR2 signaling has been implicated in neuronal injury in multiple contexts, and systemic CXCL1 administration is sufficient to induce selective dopaminergic neurons *in vivo*^62^. Here, we show that CXCL1 is elevated in conditioned media of ATP13A2-deficient astrocytes and that recombinant CXCL1 induces dose-dependent apoptosis and preferential loss of TH^+^ neurons in otherwise healthy midbrain cultures. These findings position CXCL1 as an active driver of neuronal vulnerability, and the selective sensitivity of dopaminergic neurons to intermediate concentrations of CXCL1 suggests that chemokine signaling may interact with intrinsic features of dopaminergic neurons, such as high metabolic demand and oxidative stress sensitivity^84,85^. Moreover, we find that CXCL1 levels are increased in the cerebrospinal fluid of Atp13a2^-/-^ mice, supporting the *in vivo* relevance of chemokine dysregulation as a downstream consequence of ATP13A2 loss.

Although ATP13A2 deficiency has been linked to altered α-synuclein handling, ATP13A2-associated PD does not consistently present with prominent α-synuclein pathology in patients^27–30^. In our models, we observe delayed lysosomal processing and accumulation of α-synuclein, which likely reflects a secondary consequence of lysosomal impairment rather than a primary α-synuclein-driven pathogenic mechanism. We did not assess pathogenic phosphorylated α-synuclein species, and our findings support a model in which astrocyte-mediated inflammatory signaling is the dominant driver of dopaminergic neurotoxicity in the context of ATP13A2 loss.

Finally, our identification of AMD1 as a nodal regulator connecting polyamine biosynthesis to SAM depletion highlights a therapeutically tractable pathway. Inhibition of AMD1 normalized chromatin states and prevented non-cell-autonomous dopaminergic neurotoxicity *in vitro* while attenuating astrocyte reactivity *in vivo*. While systemic targeting of polyamine metabolism will require careful consideration, these findings suggest that modulating metabolic-epigenetic coupling in astrocytes may represent a viable strategy to attenuate neuroinflammation and slow disease progression in ATP13A2-associated PD and potentially other lysosomal forms of the disease.

### Methods iPSC cultures

Human male donor-derived iPSC KOLF 2.1J lines (ATP13A2^c.1306^ and parental untargeted line, ATP13A2^WT^) were kindly donated by Bill Skarnes, iPSC Neurodegenerative Disease Initiative (iNDI)^40^.

hiPSC were maintained at 37°C, 5% CO_2_ in a humidified incubator. Cells were cultured on Geltrex-coated plates (Thermo Fisher, #A1413201) in StemFlex medium (Gibco, #A3349401). When cells reached 70-80% confluency, they were dissociated using Accutase (STEMCELL Technologies, #07920), and replated in StemFlex, supplemented during the first 24h with 10µM Rock inhibitor Y-27632 (Tocris, #1254).

### Midbrain organoids differentiation

For midbrain organoid differentiation, hiPSCs were cultured in StemFlex and differentiated as previously described^42^. Briefly, 4 x 10^7^ cells were dissociated in Accutase and placed in 125 ml spinner flasks (Corning, #3152) on a stir plate in StemFlex media supplemented with 10µM Rock inhibitor Y-27632 (Tocris, #1254). Once spheres were formed, they were filtered (pluriSelect filters) to maintain spheres ranging between 300µm and 500µm. Differentiation was then started in day 0-1 media, by dual-SMAD inhibition with SB431542 (10µM: Stemgent, #04-0010), LDN193189 (100nM: Tocris, #6053) in a base media composed of DMEM F12+Glutamax (Gibco, #10565018), supplemented with 1x B27 supplement without VitA (Gibco, #12587010), and 1X N2 supplement (Gibco, #12587010). The d0-1 media was additionally supplemented with Purmorphamine (2 μM; STEMCELL Technologies, #72202) and SAG (1 μM; Cayman Chemical, #11914) until day 3. On d4-7, midbrain patterning was achieved by adding 3µM of CHIR99021 (Tocris, #99021) to the media. Starting d12, organoids were kept in post-patterning maturation media containing the base media supplemented with 20 ng/ml GDNF (R&D Systems, #212-GD), 20 ng/mL BDNF (R&D Systems, #248-BD), 0.2 mM ascorbic acid (Fisher BioReagents, #BP351), and 0.1 mM dibutyryl cAMP (BioLog, #D009). For long-term organoid culture, after d35, organoids were switched to a mature media, composed of base media supplemented with 10ng/ml GDNF, 10ng/ml BDNF, and 0.2 mM ascorbic acid. For conditioned media experiments, organoids were moved to low-attachment 6-well plates (Thermo Scientific, #140675) on d25, at 20 organoids in 5ml of media per well.

### Midbrain neuronal culture

HiPSCs were differentiated into midbrain neurons following a previously published protocol^45^. Briefly, cells were dissociated into single cells using Accutase and plated at high density on 2x Geltrex-coated plates, with 10µM Rock inhibitor Y-27632 for the first 24h. To induce midbrain floor plate neural progenitor differentiation, cells were cultured in a base media composed of Neurobasal media supplemented with 1X Glutamax, 1X B27 supplement without VitA, and 1X N2 supplement. Media was supplemented with 250nM LDN193189 (d0-6), 10µM SB431542 (d0-6), 1µM SAG (d0-6), and different concentrations of the WNT activator CHIR (0.7µM d0-3, 7.5µM d4-9, and 3µM d10-12). Starting d10, post-patterning maturation media was used, containing the base media supplemented with 20 ng/ml GDNF (R&D Systems, #212-GD), 20 ng/mL BDNF (R&D Systems, #248-BD), 0.2 mM ascorbic acid (Fisher BioReagents, #BP351), 0.1 mM dibutyryl cAMP (BioLog, #D009), 1ng/ml TGFB3 (Peprotech, #100-21). For long-term culture, after d35, cultures were switched to a mature media, composed of base media supplemented with 10ng/ml GDNF, 10ng/ml BDNF, and 0.2 mM ascorbic acid.

### NGN2 neuronal culture

The differentiation of iPSC into neurons was performed by adapting a previously published protocol^86^. To enable doxycycline-inducible expression of NGN2 (Addgene Plasmid #209077), iPSCs were transfected with PiggyBac plasmids using Lipofectamine Stem Transfection Reagent (Thermo Fisher #STEM00003) following the manufacturer’s instructions. On day 0, dissociated iPSCs were seeded at a density of approximately 104,000 cells/cm² onto Geltrex™-coated plates in StemFlex™ media supplemented with 10 µM Y27632 and 5 µg/mL doxycycline. From day 1, the media was transitioned to Neurobasal N2B27 (comprising Neurobasal, 1x B-27, 1x N-2, 1x MEM-NEAA, 1x GlutaMax, and 1% penicillin-streptomycin) containing 10 µM SB431542, 100 nM LDN, 5 µg/mL doxycycline, and 5 µg/mL puromycin. Between days 3 and 6, daily media changes were conducted using Neurobasal N2B27 supplemented with 1 µg/mL puromycin and 5 µg/mL doxycycline. Following treatment with 0.5 µM Ara-C on day 6, cells were harvested with Accutase for co-cultures.

### Midbrain Astrocytes extraction and culture

Midbrain astrocytes were extracted from midbrain organoids starting at d100 of differentiation, as previously described^33^. Briefly, organoids were gently triturated in large chunks in TrypLE Select solution (Thermo Scientific, #12563029) using glass pipettes. Organoid chunks were then plated on 15cm dishes previously coated with 0.1% gelatin, in AM media (Astrocyte Medium, ScienCell #1801). Once cells were attached and proliferating, they were dissociated with TrypLE Select and re-plated in 0.1 gelatin pre-coated plates in AM media for a maximum of 5 extra passages. For experiments, cells were switched to an FBS-free medium. One day after passaging, the media was changed for Astrocyte Mature Media, composed by 1 part of Advanced DMEM/F-12 (ThermoFisher, #12634010), 1 part of Neurobasal (ThermoFisher, #21103049), supplemented with 1X B27 supplement without VitA, 1X N2 supplement, 1X MEM Non-Essential Amino Acids Solution (ThermoFisher, #11140050), 1X GlutaMAX (ThermoFisher, #35050061), and 10 ng/ml CNTF (PeproTech #450-13). Cell media was changed every 4 days. For cell treatments, AMD1 inhibitor MGBG (MedChem Express, HY-106634) was diluted in water at 100mM and used at a final concentration of 1µM in media for two weeks. AMXT-1501 was resuspended in DMSO at 10mM and was used in media at 5µM for 48h.

### Co-cultures

Midbrain astrocytes and neurons (either Day 25 midbrain-specific or Day 6 NGN2-induced) were dissociated using TrypLE Select and Accutase, respectively. The cells were pooled at a 5:1 (neuron-to-astrocyte) ratio and centrifuged. The resulting pellet was resuspended in cold Geltrex supplemented with 10 µM Y-27632. This suspension was then seeded as 5 µL droplets, each containing 100,000 neurons and 20,000 astrocytes, into a black 96-well plastic-bottom plate (Greiner, #). Following a 10-minute polymerization at room temperature, wells were filled with stage-appropriate neuronal differentiation media supplemented with 10 ng/mL CNTF. Cultures were maintained for four weeks via 50% media exchanges every 48 hours.

### Astrocyte Conditioned Media Experiments

To generate astrocyte-conditioned media (ACM), astrocytes were maintained in Mature Astrocyte Media, with the supernatant harvested four days post-feeding. The conditioned media were centrifuged at 2,000 x g for 5 minutes to remove cellular debris and subsequently stored at -80°C. D25 midbrain organoids or 2D neuronal cultures were then maintained in the conditioned media for 30 days, with 50% media exchanges performed every three days.

### CXCL1 supplementation on midbrain neurons

To assess the effects of CXCL1 on midbrain neurons, conditioned media collected from ATP13A2^WT^ astrocyte cultures were supplemented with recombinant CXCL1 (PeproTech, 275-GR-010) at concentrations as indicated in each experiment. Recombinant CXCL1 was diluted in 0.1% BSA in PBS and then in conditioned media. Primary midbrain neuronal cultures were exposed to the supplemented conditioned media for the four weeks under standard culture conditions. Control conditions consisted of ATP13A2^WT^ astrocyte conditioned media without added CXCL1.

### Animals and genotyping

Mice were maintained under a 12 h light/12 h dark cycle with ad libitum access to food and water. All experimental procedures were approved by the Bioethical Committee of KU Leuven and conducted in accordance with institutional guidelines (Project number P069-2021) and conformed to the guidelines of the European Communities Council Directive of November 24, 1986 (86/609/EEC).

*Atp13a2* wild-type (WT) and knockout (KO) mice were generated by intercrossing *Atp13a2* heterozygous animals on a C57BL/6J background. Heterozygous mice were produced by crossing male *Atp13a2*-flox mice (LoxP sites flanking exons 2-3; RRID: IMSR_JAX:028387) with female Sox2-Cre transgenic mice (RRID: IMSR_JAX:008454) to achieve germline excision of the floxed Atp13a2 allele.

Genotyping was performed on genomic DNA extracted from tail biopsies collected at postnatal day 2. Polymerase chain reaction (PCR) was conducted using two primer sets: Forward 5′-CTG CAG CTT CGA GAG GAA AG-3′; Reverse (WT/Floxed) 5′-CAC TCT GTC CTC AGG CCT TC-3′; Reverse (KO) 5′-TGA GAA GTG GGA ATC GGG-3′. PCR products were resolved on 1.5% agarose gels stained with Midori Green. Amplicons of 426 bp and 256 bp corresponded to the WT and KO alleles, respectively.

MGBG was administered at 2mg/kg through weekly subcutaneous injections in *Atp13a2* WT and KO starting at 2 weeks until 10 weeks of age {Seppänen, 1983 #68; Amlacher, 1990 #69}. Saline was administered as control.

### Protein Extraction and Immunoblotting

Cells were lysed in RIPA buffer (Thermo Scientific, #89900), supplemented with protease and phosphatase inhibitor cocktails (MilliporeSigma, #11836153001 and Abcam, #ab201113), or a 2% SDS lysis buffer (10 mM Tris, 150 mM NaCl, 1 mM EDTA, 2% SDS). Protein concentrations were determined using a Pierce BCA Protein Assay Kit (Thermo Scientific, #23225). The sample mix was prepared in 1x Laemmli Buffer (BioRad, #1610747) and denatured for 5 minutes at 95 °C. Samples were loaded onto XT Precast gel (Bio-Rad, #340123), then transferred to a PVDF membrane (0.22 µm; Bio-Rad, #1620177) using a semi-dry blotting system (Bio-Rad, #1704150). Membranes were blocked in 5% BSA in TBS-T (TBS 1x, 0.1% v/v Tween 20) for 1h at RT, and incubated with primary antibodies (Table 1) overnight on a shaker at 4 °C. Membranes were then incubated in the appropriate secondary HRP-conjugated antibody for 1h at RT and imaged using ECL (PerkinElmer, #NELI03001EA) on a Licor imager (Odyssey® XF Imaging System). For dot blot analysis, 3µg of quantified protein from collected media was applied directly on dry nitrocellulose membranes in small drops; the membranes were then blocked and immunoblotted as previously described in the paragraph. All densitometric quantifications were performed using Fiji software.

**Table 1:** Primary antibody list

| Antibody | Source | Identifier |
| --- | --- | --- |
| α-synuclein | Abcam | #AB138501 |
| Actin | EMD Millipore | #S2532 |
| Aggregated-α-synuclein | Abcam | #AB209538 |
| ATP13A2 | Sigma | #A3361 |
| CD44 | Cell Signaling Technology | #3570S |
| DAPI | Cayman Chemical | #13197 |
| FOXA2 | Abcam | #AB60721 |
| GAPDH | Abcam | #ab9485 |
| GFAP | EMD millipore | #AB5804 |
| GFAP | Dako | #AB_10013382 |
| GIRK2 | Alomone labs | #APC-006 |
| Histone 3 K9 dimethylation | Abcam | #AB_449854 |
| Histone 3 K4 trimethylation | Abcam | #AB_306649 |
| LAMP1 | Novus biotech | #NBP2-25183 |
| MAP2 | BioLegend | #822501 |
| Recombinant α-synuclein-HiLyte Fluor 488 labeled | Anaspec | #AS-55457 |
| S100A6 | Abcam | #ab181975 |
| S100B | EMD millipore | #S2532 |
| SOX9 | Abcam | #AB185966 |
| TH | Abcam | #ab112 |
| TUJ1 | BioLegend | #801202 |

### Lentiviral Production and Transduction

Lentivirus particles were generated in-house with HEK293T cells maintained in DMEM medium supplemented with 10% bovine calf serum. Cells were transfected using the calcium phosphate method, utilizing a mixture of the transfer vector, three packaging plasmids (pMDL, pRev, and pVSVG), CaCl_2_, and BBS. The supernatant was harvested at 48 and 72 hours post-transfection, cleared of cellular debris via centrifugation (2,000 g, 10 minutes, 4 °C), and passed through a 0.45µm filter. To concentrate the viral particles, the supernatant was incubated with 5% PEG 8000 and 3% 5M NaCl overnight at 4 °C, followed by centrifugation at 3000 g for 20 min. at 4°C. The resulting viral pellet was resuspended in DMEM/F12 + GlutaMAX and stored as single -use aliquots at -80°C. Viral titers were determined in HEK293T cells; the lowest concentration yielding transduction efficiency was utilized for subsequent experiments. For astrocyte transduction, the virus was added to the culture media at the time of plating. The vectors employed included pFUw-LAMP1Sig-mCherry (Addgene #239556)^61^ and Sigma MISSION shRNA vectors: shScrambled (#SHC202), shAMD1#1 (GATGGAACTTATTGGACTATT), and shAMD1#2 (GTCTCCAAGAGACGTTTCATT). Following infection, stable astrocytes were selected with puromycin for 3 days before downstream analysis. The polyamine sensor plasmid (Addgene #232356)^64^ was kindly provided by Dr. Ankur Jain.

### Live-Cell imaging of α-synuclein internalization

Astrocytes were transduced with mCherry-LAMP1 lentivirus^61^ four days before replating for imaging to allow for stable expression of the lysosomal marker. On the day of the experiment, cells were incubated with 2µg/ml HiLyte™ 488-labeled human recombinant α-synuclein (AnaSpec, #AS-55457), and with 10µg/ml Hoechst 33342. Live-cell imaging was performed at the indicated time points.

### RNA Extraction and RT-qPCR

Total RNA was extracted using the Monarch Total RNA Miniprep Kit (New England BioLabs, #T2010S). cDNA synthesis was performed on 1 μg of RNA using the High-Capacity cDNA Reverse Transcription Kit (Applied Biosystems, #4368814). For quantitative PCR (qPCR) analysis, cDNA was diluted 1:10 and combined with PowerUp SYBR Green Master Mix (Applied Biosystems, #A25742) and gene-specific primers (Table 2). Reactions were performed using a QuantStudio 7 Flex Real-Time PCR System (Applied Biosystems, #4485701). Relative gene expression was calculated using the 2^-ΔΔCT^ method, with *ACTB* used for normalization.

**Table 2.**
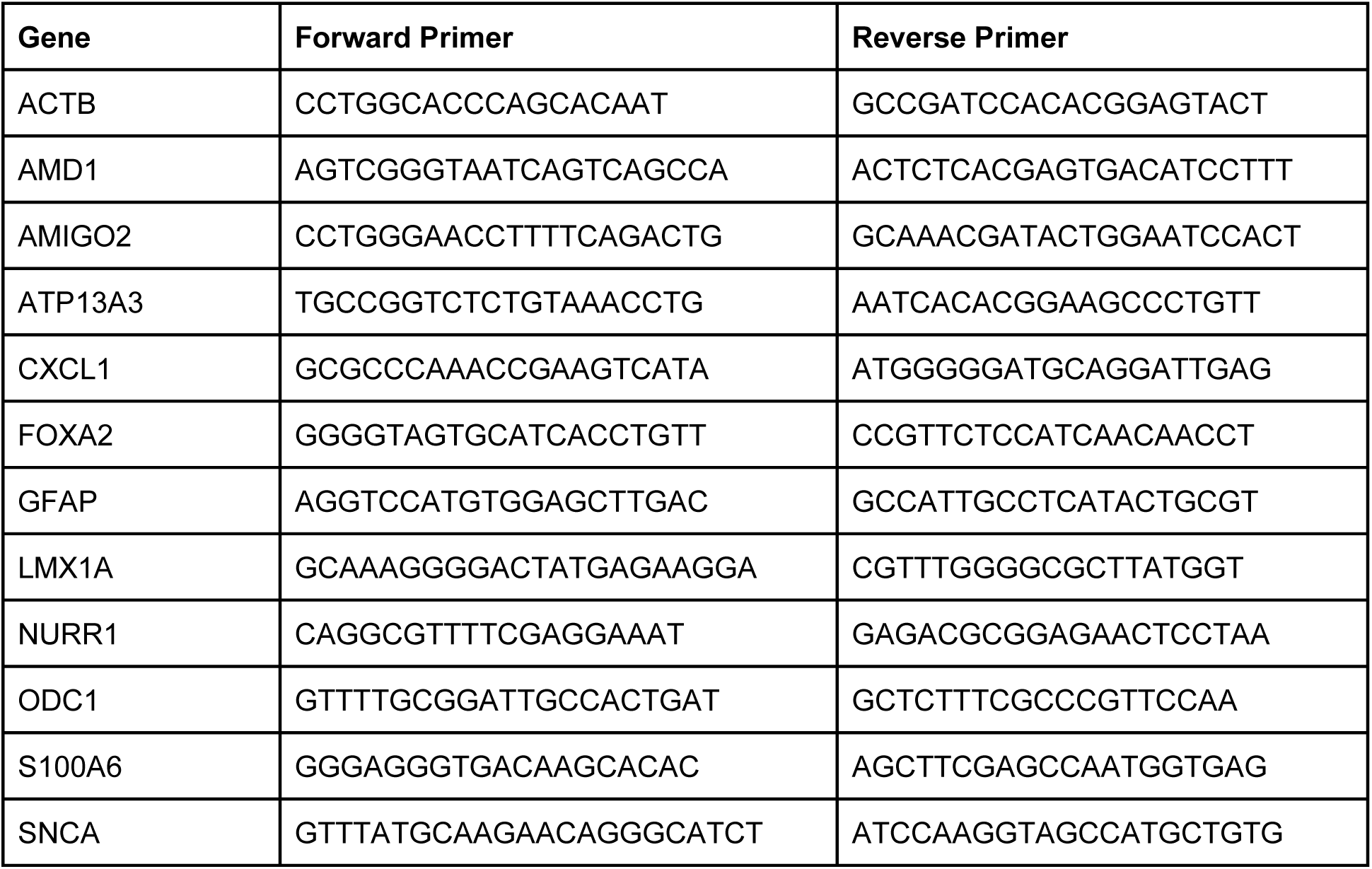
– Primer list

### Bulk RN2A-sequencing

Total RNA was extracted using the Monarch Total RNA Miniprep Kit (New England BioLabs, #T2010S). Sample quality control, library preparation, and standard RNA-sequencing were performed by GENEWIZ LLC (South Plainfield, NJ). FASTQ files were pseudo-aligned to the reference genome (GRCh38.p13) using the *kallisto* pipeline to obtain transcript abundance estimates. Differential expression analysis was subsequently conducted using the *edgeR* package in R (version 2023.12.0+369). Data visualization and functional enrichment analyses were performed using the *ggplot2* and *clusterProfiler* packages.

### Bulk ATAC-sequencing

Cell pellets were collected and submitted to GENEWIZ LLC (South Plainfield, NJ) for ATAC-sequencing (Assay for Transposase-Accessible Chromatin). To identify differentially accessible regions (DARs), raw reads were aligned to the human reference genome (GRCh38.p13), and peak calling was performed. Differentially open peaks were identified and analyzed to determine chromatin accessibility across experimental conditions. Data visualization and functional enrichment analyses were performed using the *ggplot2* packages in R (version 2023.12.0+369). Peak visualization was generated in IGV.

### DNA extraction and Bisulfite sequencing

Genomic DNA was isolated from astrocytes using the QIAamp DNA Mini Kit (Qiagen) following the manufacturer’s protocol. Shortly, astrocytes were harvested using 1 mL of TrypLE E in one well of a 6-well plate. Cells were then centrifuged and processed for column-based DNA purification. DNA was eluted in a final volume of 80uL and quantified before downstream processing, yielding concentrations ranging from 51 to 198 ng/uL. Purified DNA was subsequently submitted to Zymo Research for Reduced Representation Bisulfite Sequencing (RRBS). RRBS reads were processed with CutAdapt, aligned with Bismark (hg38), and analyzed in R using methylKit. Bisulfite conversion efficiency exceeded 99% in all samples. Differential methylation was assessed using logistic regression with FDR correction, and differentially methylated regions were annotated using UCSC genome annotations.

### Immunohistochemistry and Histological Analysis

Organoids were washed with PBS without calcium and magnesium (PBS^-/-^) and fixed in 4% paraformaldehyde (PFA) overnight at 4°C with gentle agitation. Following fixation, samples were dehydrated and embedded in paraffin. Sectioning, staining, and subsequent imaging were performed by the Neuropathology Brain Bank and Research Core (NPBB) at the Icahn School of Medicine at Mount Sinai.

For mice brain samples, 10-week-old *Atp13a2* knock out (-/-) and wild type (+/+) mice were subjected to transcardial perfusion for tissue collection. Animals received a lethal intraperitoneal dose of sodium pentobarbital (100 mg/kg; Dolithol), followed by perfusion with 0.9% saline and subsequently 4% paraformaldehyde (PFA; Sigma-Aldrich, 441244). Brains were dissected and post-fixed overnight in 4% PFA. Sagittal sections (40 µm) were obtained using a vibrating microtome and processed for immunohistochemistry.

Free-floating sections were washed in PBS and incubated for 40 min in blocking solution containing 10% donkey serum (Jackson ImmunoResearch, 017-000-121, RRID: AB2337258). Sections were incubated overnight at room temperature with rabbit anti-CD68 (1:500; Abcam, ab125212, RRID: AB_10975465) diluted in blocking solution. After PBS washes, sections were incubated for 2 h at room temperature with donkey anti-rabbit Alexa Fluor 488 (1:500; Abcam, ab150073, RRID: AB_2636877). Sections were mounted onto pre-coated slides and counterstained with DAPI (1:500; MedChemExpress, HY-D0814).

### Immunofluorescence and Confocal Microscopy

2D cultured cells, cells were seeded in 96-well black-walled plates (Greiner, #655090) and fixed with 4% PFA for 15 minutes at room temperature (RT). Permeabilization and blocking were performed for 1 hour at RT in a buffer consisting of 0.3% Triton X-100 and 5% donkey serum in PBS. Primary antibodies (Table 1) were diluted in blocking buffer and incubated overnight at 4 °C. Following three PBS washes, wells were incubated with species-specific Alexa Fluor-conjugated secondary antibodies for 1 hour at RT. Nuclei were counterstained with Hoechst 33342 (Cayman Chemical, #13197) for 5 minutes.

Organoids were harvested, washed with PBS without calcium or magnesium (PBS-/-), and fixed in 4% PFA overnight at 4 °C with gentle agitation. For cryopreservation, organoids were incubated in 30% sucrose overnight at 4 °C, subsequently embedded in OCT (Tissue-Tek, #4583), and cryosectioned at 30 µm thickness. Sections were mounted on Superfrost Plus Microscope Slides (Fisher Scientific, #12-550-15) and processed for immunofluorescence using the primary antibodies and blocking conditions described above.

### Effector Caspase assay

To evaluate apoptosis, active caspase-3/7 was visualized using the Image-iT ™ LIVE/DEAD™ Caspase-3/7 Detection Kit (Thermo Fisher Scientific) following the manufacturer’s protocol. iPSC-derived neurons were incubated with the detection reagent for 1 h at 37 °C. Immediately following the live-cell labeling, the cultures were fixed in 4% paraformaldehyde for 15 min at room temperature. Subsequently, the fixed samples were processed for immunofluorescence analysis.

### Microscopy and Image Acquisition

Confocal immunofluorescence images were acquired using a Nikon Ti2-E AX Confocal Microscope or a High Content Imager CX7 (Thermo Fisher) with a HCS Studio 4.0. For histological analysis, immunohistochemistry (IHC) slides were scanned at 40x magnification using an Aperio VERSA 8 digital slide scanner (Leica Biosystems, Wetzlar, Germany). Representative fields of interest were identified and captured using Aperio ImageScope Software (Leica Biosystems). To ensure consistency, all images within a given experiment were acquired using identical laser power, gain, and exposure settings.

For brain samples imaging, GFAP-immunolabelled sections were imaged with LSM880 Airyscan confocal at 40x magnification, z-stack images were captured at a focal density of 0.43 µm with a pinhole of 1 AU, scan speed of 0.5 and averaging of 2

### Image Analysis and Quantification

Digital image processing and figure preparation were performed using Fiji/ImageJ software. For quantitative immunofluorescence analysis, CellProfiler software was utilized to develop automated pipelines for cell counting, area-based thresholding, colocalization analysis and signal intensity measurements. Specifically, the software was used to identify nuclei and cellular boundaries to determine the number of cells positive for specific markers and the total area occupied by staining.

For the analysis of immunohistochemistry (IHC) in sectioned organoids, QuPath software was employed. Automated cell detection and classification algorithms were utilized to quantify marker expression across whole-slide images, ensuring unbiased sampling of the organoid tissue.

### LDH Cytotoxicity Assay

To quantify cell death and cytotoxicity, the LDG-Glo^TM^ Cytotoxicity Assay (Promega, #J2380) was employed. Briefly, 2µl of conditioned media was harvested and diluted in 298µl of storage buffer (200 mM Tris-HCl pH 7.3, 10% glycerol, 1% BSA) and stored at -20 °C before analysis. Once all experimental time points were collected, the assay was performed according to the manufacturer’s instructions. Samples were mixed with the detection reagent (1:1 ratio) and incubated at room temperature for 1h. Luminescence was measured using a Varioskan Luxmicroplate reader (Thermo Fisher).

### Lysosomal Function and Phagocytosis Assays

To assess lysosomal pH, cells were incubated with 1 μM LysoSensor™ Green DND-189 (Thermo Fisher Scientific, #L7535) for 1 minute at 37 °C and immediately processed for flow cytometry. For the evaluation of hydrolytic lysosomal capacity, cells were pulse-labeled with 1 μg/mL DQ™ Red BSA (Thermo Fisher Scientific, #D12051) for 30 minutes at 37 °C. Following a media exchange and a 6-hour incubation to allow for protein degradation and fluorophore release, cells were dissociated and prepared for analysis.

Briefly, cells were pelleted and resuspended in PBS supplemented with 2% BSA and filtered through a 35 μm cell strainer. DAPI (Cayman Chemical, #13197) was added to exclude dead cells from the analysis. Fluorescence Minus One (FMO) controls were utilized for gating. Single-cell data were acquired using a BD FACS Celesta™ flow cytometer and analyzed using FCS Express 7 software (De Novo Software).

Lysosomal proteolytic activity was further monitored over time by imaging DQ-BSA-labeled cells every hour for 20 hours using the Incucyte® live-cell imaging system. Integrated intensity was calculated and normalized to well confluence using the Incucyte® analysis software.

For phagocytosis assessment, purified myelin was labeled with pHrodo™ Red according to the manufacturer’s instructions. Cells were exposed to pHrodo-labeled myelin, and fluorescence was acquired hourly over 24 hours. The total red fluorescence intensity, representing myelin internalization into acidic phagosomal compartments, was quantified and normalized to cell confluence.

### Cytokine Profiling

To evaluate the inflammatory secretome of astrocytes, conditioned media were analyzed using the Proteome Profiler™ Human XL Cytokine Array Kit (R&D Systems, #ARY022B) according to the manufacturer’s instructions.

Following the assay, the membranes were developed using chemiluminescence and imaged. The integrated optical density of each spot was quantified using Fiji software. Background subtraction was performed, and the signal intensities were normalized to the internal positive control spots on each membrane to determine relative protein expression levels across experimental groups.

### Polyamine Uptake Assay

To quantify the internalization of specific polyamines, BODIPY-spermidine and BODIPY-spermine (synthesized and kindly provided by the laboratory of Dr. Peter Vangheluwe) were utilized. The fluorescent polyamine probes were added to the culture media at a final concentration of 5 μM. Following a 4-hour incubation at 37 °C, the cells were washed with PBS to remove non-internalized probes and dissociated.

Cells were processed for flow cytometry analysis as described in the Lysosomal Function and Phagocytosis Assays methods. Briefly, cells were resuspended in PBS supplemented with 2% BSA, filtered through a 35 μm strainer, and stained with DAPI (#13197) to exclude dead cells. Single-cell fluorescence intensity was acquired using a BD FACS Celesta™ flow cytometer and analyzed using FCS Express 7 software (De Novo Software) to determine the mean fluorescent intensity (MFI) of polyamine uptake.

### S-Adenosylmethionine (SAM) Quantification

Astrocytes were harvested by scraping in ice-cold PBS and pelleted by centrifugation at 1000 x g for 5 minutes at 4 °C. S-Adenosylmethionine (SAM) levels were quantified using a competitive ELISA kit (Cell Biolabs Inc., #MET-5152) according to the manufacturer’s instructions.

Before the assay, total protein concentration was determined using a BCA Protein Assay to ensure equal loading and for final normalization of SAM concentrations. Absorbance was measured at 450 nm using a Varioskan Lux microplate reader (Thermo Fisher). The SAM concentration in each sample was calculated against a standard curve and expressed as nmol/mg of total protein.

### CXCL1 Elisa

CXCL1 levels were quantified using a commercially available ELISA kit (Abcam, #AB190805) according to the manufacturer’s instructions. Conditioned media were collected from astrocyte cultures, centrifuged to remove debris, and analyzed without dilution. Standards and samples were loaded in duplicate onto pre-coated 96-well plates and incubated with the provided detection reagents following the kit protocol. After the final wash steps, signal was developed using the supplied substrate solution and stopped with stop buffer. Absorbance was measured at 450 nm using Varioskan Lux microplate reader (Thermo Fisher), and CXCL1 concentrations were calculated from a standard curve generated with recombinant CXCL1 standards included in the kit.

### Cerebrospinal fluid collection

Cerebrospinal fluid (CSF) was obtained from 10-week-old *Atp13a2* knockout (Atp13a2-/-) and wild type (Atp13a2+/+) mice by puncture of the cisterna magna. Animals were anesthetized and positioned in a stereotaxic apparatus. After shaving and disinfecting the dorsal cervical region, neck muscles were gently separated to visualize the cisterna magna. A fine glass capillary mounted on a micromanipulator was advanced at approximately 45° angle while avoiding blood vessels, allowing CSF to be drawn by capillary action. Samples were centrifuged at 10,000 x g for 10 min at 4 °C to remove particulate material and stored at -80 °C.

### CXCL1 quantification in CSF

CXCL1 levels were measured in CSF samples using a V-PLEX Proinflammatory Panel 1 Mouse Kit (K15048D, Meso Scale Discovery) according to the manufacturer’s protocol. Briefly, samples and calibration standards were added to plates coated with analyte-specific capture antibodies. Bound proteins were detected with SULFO-TAG–labeled secondary antibodies, and electrochemiluminescence signals were acquired using a MESO QuickPlex SQ 120MM reader (Meso Scale Discovery).

## Statistical Analysis

All experiments were performed in at least three independent biological replicates. Data are presented as the mean±standard error of the mean (SEM).

Statistical evaluations and graphical representations were generated using GraphPad Prism software (version 10.6.0). Comparisons between two groups were evaluated using a paired or unpaired two-tailed Student’s *t*-test, as appropriate. For experiments involving more than two groups or multiple factors, one-way or two-way analysis of variance (ANOVA) was applied, followed by Holm-Šídák multiple comparisons test to adjust for multiple testing. Statistical significance was defined as p<0.05.

## Availability Statement

The data, protocols, and key lab materials used and generated in this study are listed in a Key Resource Table alongside their identifiers at *10.5281/zenodo.18704033*.

## Supporting information

Key Resource Table

## Acknowledgments

The authors thank Bill Skarnes, iPSC Neurodegenerative Disease Initiative (iNDI), the Center for Alzheimer’s and Related Dementias (CARD) and the ASAP consortium for the donation of the KOLF2.1J cell lines. Dr. Ankur Jain for the donation of the polyamine reporter plasmids. The Neuropathology Brain Bank & Research CoRE at the Icahn School of Medicine at Mount Sinai for the technical help in sectioning, staining, and imaging organoids.

This research was funded by Aligning Science Across Parkinson’s [ASAP-024297] through the Michael J. Fox Foundation for Parkinson’s Research (MJFF) and by NIH-NINDS [R01NS114239]. For open access, the author has applied a CC BY public copyright license to all Author Accepted Manuscripts arising from this submission .

## Declaration of generative AI and AI-assisted technologies in the manuscript preparation process

During the preparation of this work, the authors used ChatGPT (OpenAI) to assist with troubleshooting computational code and improving the clarity and grammar of the manuscript. All outputs were critically reviewed and edited by the authors, who take full responsibility for the final content of the published article.

---

**Figure S1.**
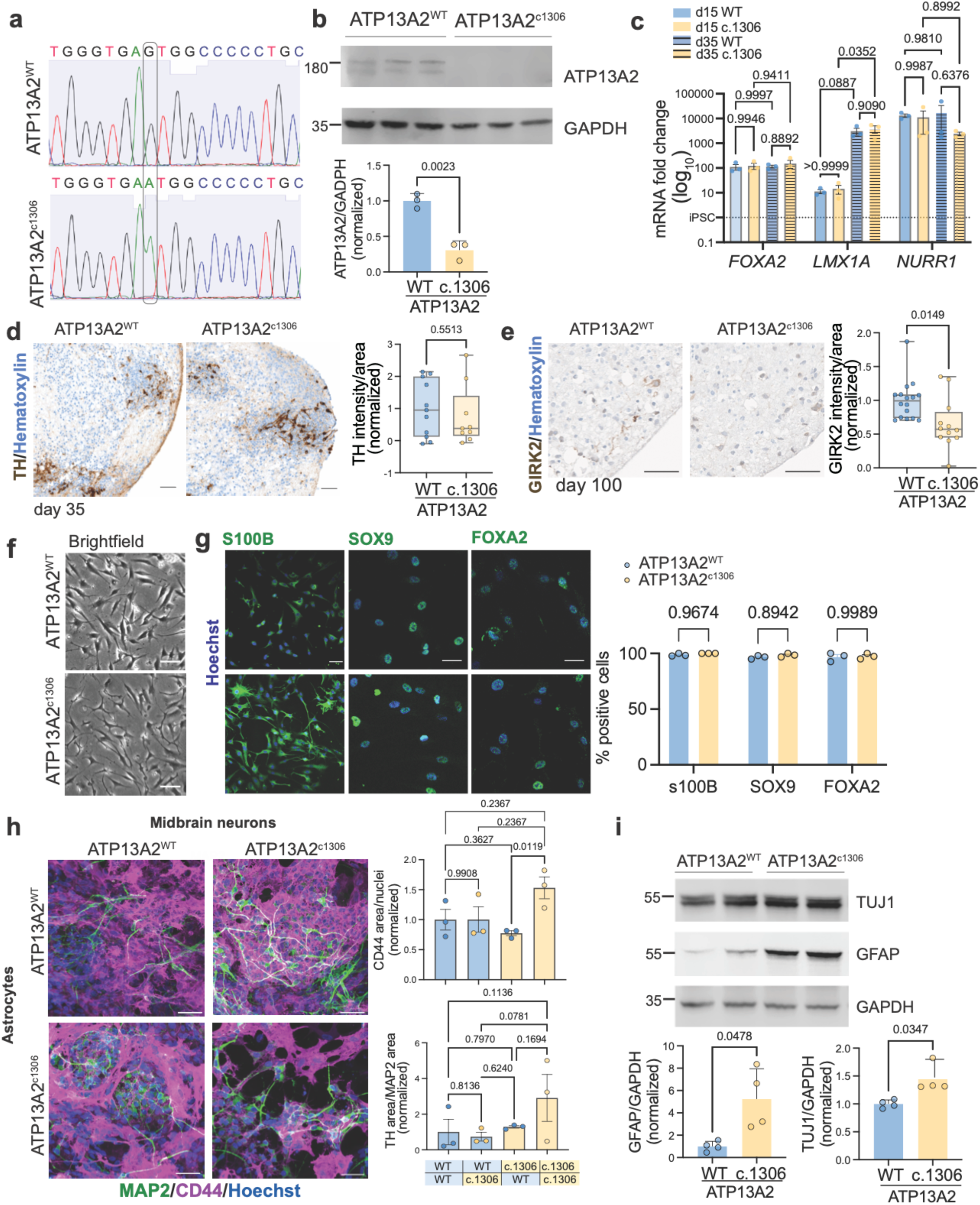
a. Sanger sequencing results of KOLF2.1J lines used for the project, which evidenced the single SNP G>A that causes the c.1306 mutation. **b.** Western blot of midbrain astrocytes of ATP13A2 and GAPDH as loading control and quantification of normalized levels of ATP13A2; differences analyzed by Two-tailed t-test. **c.** mRNA fold change by qPCR of midbrain organoids at d15 and d35 to corroborate expression of midbrain-specific markers. Undifferentiated iPSC value indicated by the undashed line (value=1). **d.** Micrography of TH (brown) and Hematoxylin (blue) of midbrain organoids at d25. Scale bar 20µm. Differences were analyzed by a two-tailed t-test. **e.** Micrography GIRK2 (brown) and Hematoxylin (blue) staining in ATP13A2^WT^ and ATP13A2^c.1306^ day 100 midbrain organoids and quantification. Scale bar 20µm. Differences were analyzed by a two-tailed t-test. **f.** Brightfield images of midbrain astrocytes in 2D culture. **g.** Immunocytochemistry of astrocytes stained for cell type (s100B, SOX9), and region (FOXA2) specific markers expression (green) and Hoechst (blue). Quantification of marker-positive cell percentage. Scale bar 50µm. **h.** Immunocytochemistry of mixed genetic co-cultures of midbrain neurons stained for MAP2 (green), midbrain astrocytes CD44 (magenta), and Hoechst (blue), and their quantification. Scale 50µm. **i.** Immunoblots of WT cortical neurons (NGN2) co-cultured with astrocytes for TUJ1 and GFAP. GAPDH was used as a loading control. Differences were analyzed by a two-tailed t-test.

**Figure S2.**
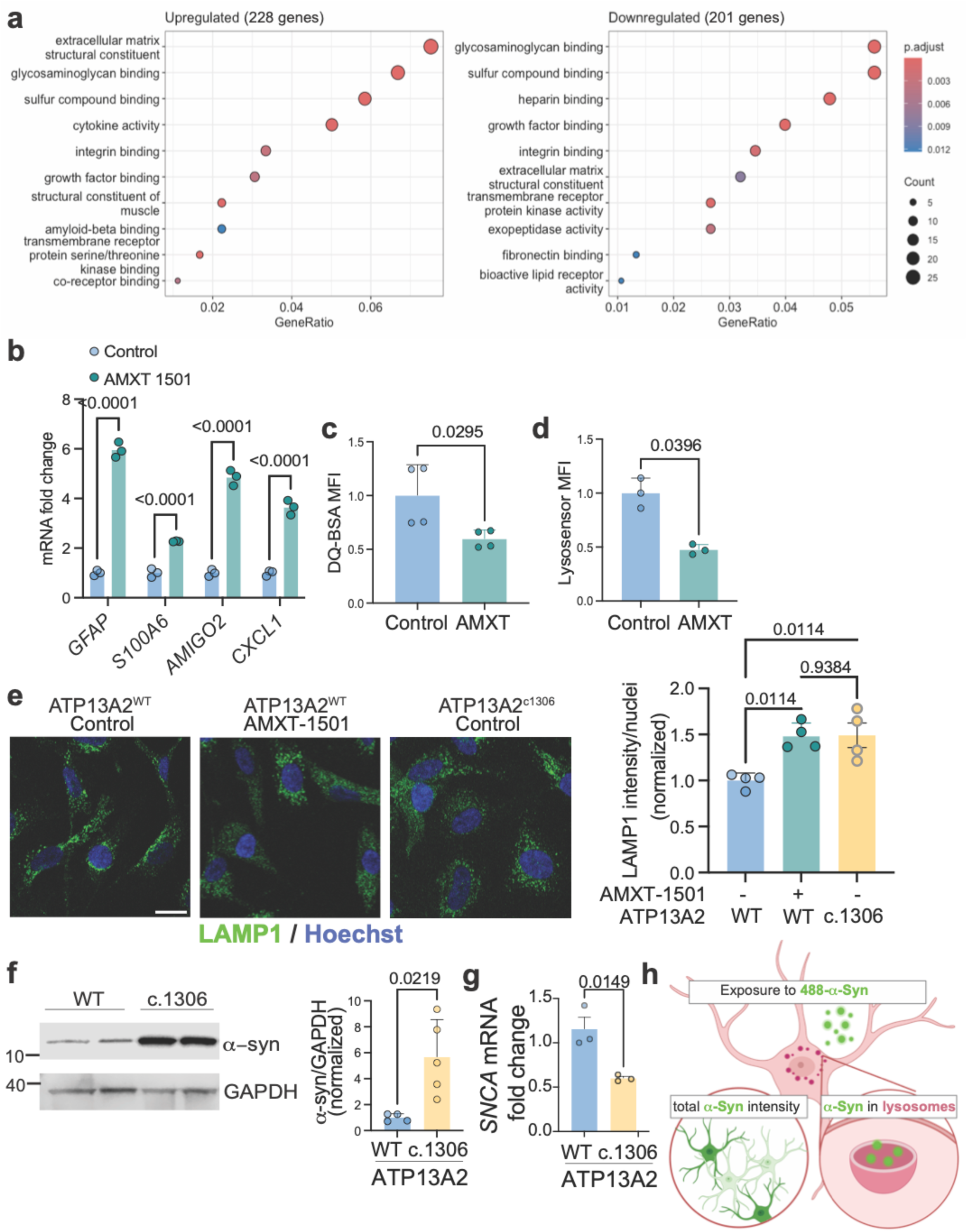
a. ClusterProfiler dot plots of the GO terms for the ATP13A2^c.1306^ vs ATP13A2^WT^ upregulated and downregulated genes detected in bulk RNAseq. **b.** mRNA fold change by qPCR of neuroinflammation-associated genes in ATP13A2^WT^ astrocytes treated with the PA transport inhibitor AMXT-1501, or control (DMSO). Differences were analyzed by Two-way ANOVA. **c.** DQBSA MFI and **d.** Lysosensor MFI of astrocytes treated with the PA transport inhibitor AMXT-1501, or control (DMSO). Differences were analyzed by a two-tailed t-test. **f.** Immunoblots of ATP13A2^WT^ and ATP13A2^c.1306^ astrocytes for α-synuclein. GAPDH was used as a loading control. Differences were analyzed by a two-tailed t-test. **g.** mRNA fold change by qPCR of *SNCA*. differences analyzed by Two-tailed t-test **h.** Schematic of α-synuclein uptake experiments. Astrocytes transduced with mCherry-LAMP1 viruses were exposed to fluorescently labeled α-synuclein monomers. Live imaging over time allowed us to quantify overall total α-synuclein in cells and co-localizations with LAMP1-positive signal (lysosomes).

**Figure S3.**
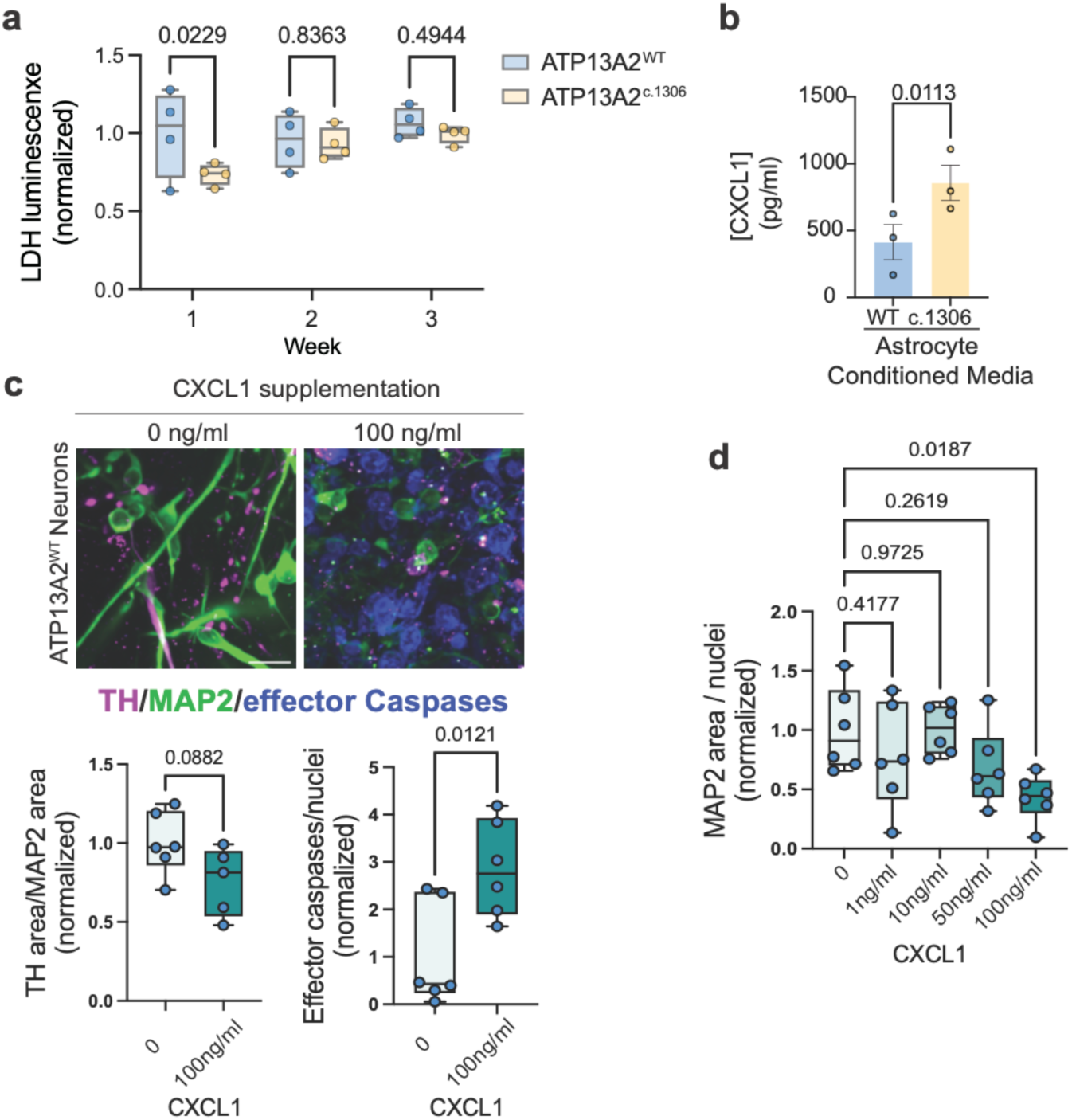
a. LDH assay in ATP13A2^WT^ exposed to astrocyte condition media at week one to three. **b.** ELISA for CXCL1 in conditioned media from astrocytes, differences analyzed by paired Two-tailed t-test. **c.** Immunofluorescence of TH (magenta), MAP2 (green), and active (cleaved) effector caspase 3/7 (blue) in neurons exposed to ATP13A2^WT^ conditioned media supplemented with 100ng/ml of recombinant CXCL1. Differences were analyzed by a two-tailed t-test. **d.** Quantification of MAP2/nuclei in neurons exposed to ATP13A2^WT^ conditioned media supplemented with increasing amounts of recombinant CXCL1 (Fig 3h and S3c). Differences were analyzed by One-way ANOVA.

**Figure S4.**
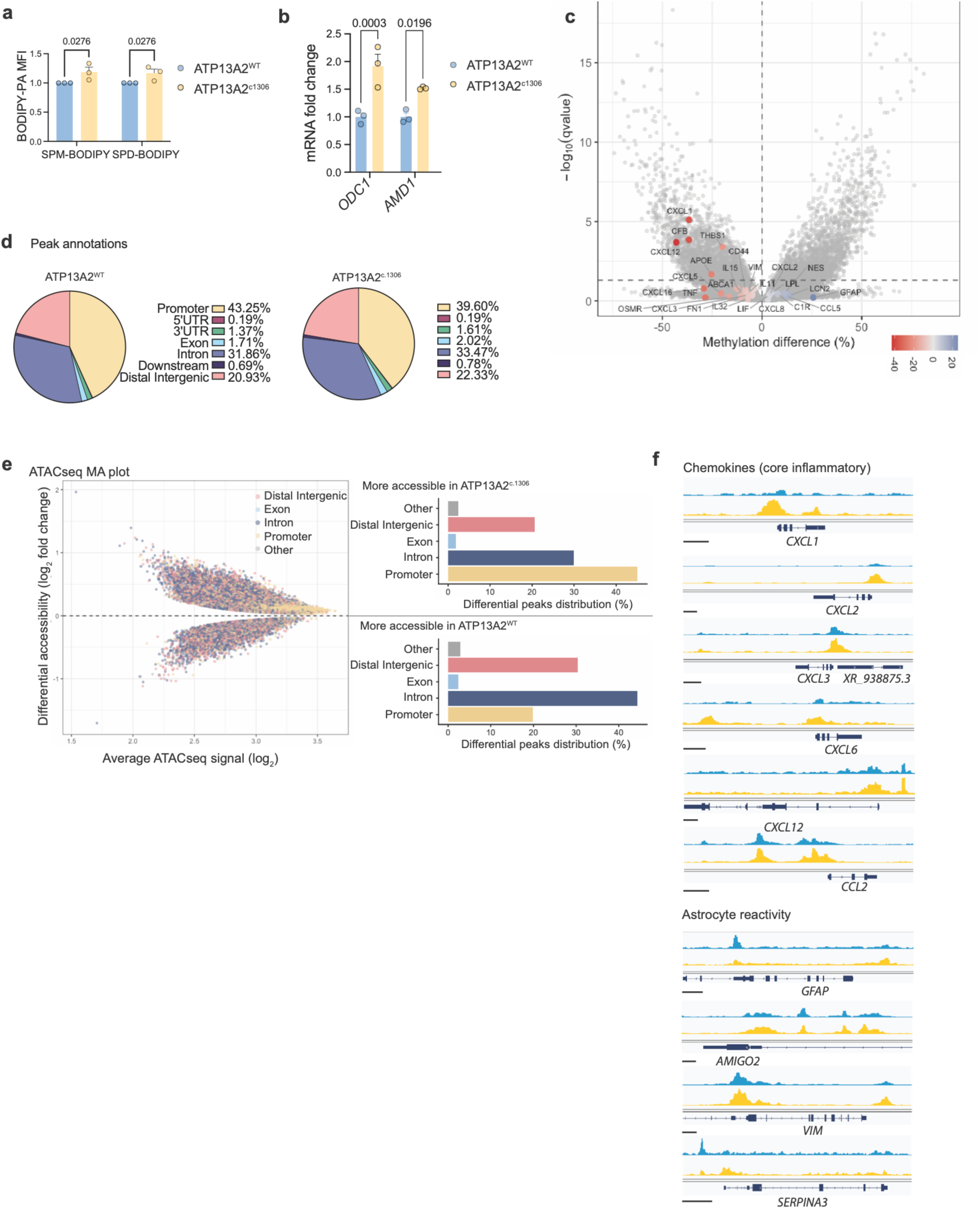
a. Quantification of BODIPY-labeled and spermine (SPM) and spermidine (SPD) in astrocytes through median fluorescence intensity. Differences were analyzed by Two-way ANOVA. **b.** mRNA fold change by qPCR of *ODC1* and *AMD1*. Differences were analyzed by Two-way ANOVA. **c.** Volcano plot of differentially methylated promoters in ATP13A2^c1306^ vs ATP13A2^WT^ astrocytes. Labeled neuroinflammatory genes color-coded per methylation difference % **d.** Pie chart showing distribution of the differentially open peaks detected by ATAC-seq in ATP13A2^WT^ and ATP13A2^c1306^ astrocytes **e.** MA plot displaying peak accessibility in ATP13A2^c1306^ vs. ATP13A2^WT^ astrocytes. M (y-axis) denotes the log2fold change and A (x-axis) the mean of normalized counts. Individual peaks are color-coded by their annotated genomic regions. Bar plots summarize the percentage of DARs localized to specific genomic features **f.** Genomic snapshots of selected inflammatory signaling and astrocyte reactivity genes in ATP13A2^WT^ (blue) and ATP13A2^c1306^ (yellow) astrocytes. Tracks show normalized peak heights.

**Figure S5.**
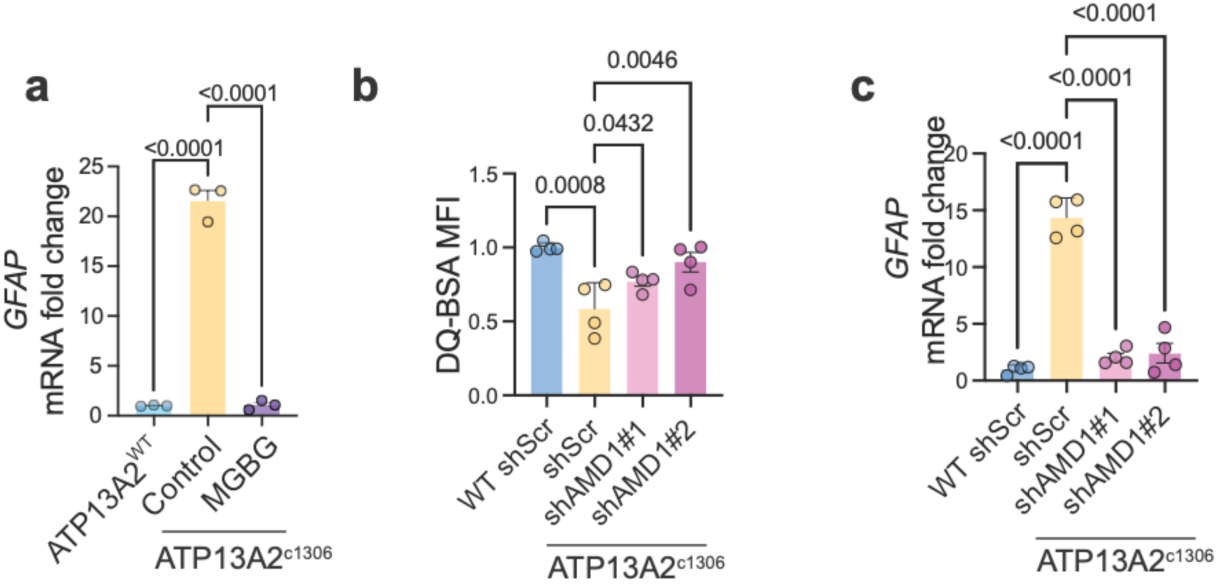
a. mRNA fold change by qPCR of *GFAP* in ATP13A2^WT^ and ATP13A2^c.1306^ astrocytes treated with MGBG or control solution (H2O). Differences were analyzed by One-way ANOVA. **b.** Median fluorescence intensity of DQ-BSA to measure proteolytic function in astrocytes targeted with shRNA for AMD1 or control shScramble. Differences were analyzed by One-way ANOVA. **h.** mRNA fold change by qPCR of *GFAP* in astrocytes targeted with shRNA for AMD1 shScramble. Differences were analyzed by One-way ANOVA.

