## Supplementary material for "ATP13A2 Loss of Function-Driven Polyamine Dysregulation Induces SAM Depletion and Epigenetic Astrocyte Toxicity": Key Resource Table

| Resource Type | Resource Name | Source | Identifier | New/Reuse | Additional info |
|---|---|---|---|---|---|
| **Dataset** | bulkRNA-seq data from WT, ATP13A2 c.1306 | This paper | https://doi.org/10.5281/zenodo.19226018 | new | |
| **Dataset** | bulkATAC-seq data from WT, ATP13A2 c.1306 | This paper | https://doi.org/10.5281/zenodo.19226012 | new | |
| **Dataset** | bisulfite sequencing | This paper | https://doi.org/10.5281/zenodo.19226020 | new | |
| **code** | bulkRNA-seq data from WT, ATP13A2 c.1306 | This paper | https://github.com/blanchardlab/ATP13A2-paper/tree/main/RNAseq | new | |
| **code** | bulkATAC-seq data from WT, ATP13A2 c.1306 | This paper | https://github.com/blanchardlab/ATP13A2-paper/tree/main/ATACseq | new | |
| **code** | bisulfite sequencing | This paper | https://github.com/blanchardlab/ATP13A2-paper/tree/main/Bisulphateseq | new | |
| **Experimental model: Organism/strain** | Sox2-cre expressing female mice | This paper | RRID:IMSR_JAX:008454 | reuse | |
| **Experimental model: Organism/strain** | Atp13a2 Flox male mice | This paper | RRID:IMSR_JAX:028387 | reuse | |
| **Experimental model: iPSC** | KOLF2.1 J (ATP13A2WT) | Jax Laboratory | RRID:CVCL_B5P3 | reuse | |
| **Experimental model: iPSC** | KOLF2.1J c1306 SNV/SNV | Jax Laboratory | RRID:CVCL_F2AU | reuse | |
| **Protocol** | Brain tissue collection | protocols.io | dx.doi.org/10.17504/protocols.io.261gedq8ov47/v1 | reuse | |
| **Protocol** | Perfusion | protocols.io | dx.doi.org/10.17504/protocols.io.5jyl8p3qrg2w/v1 | reuse | |
| **Protocol** | Brain sectioning using vibratome | protocols.io | dx.doi.org/10.17504/protocols.io.j8nlko72xv5r/v1 | reuse | |
| **Protocol** | Immunofluorescent staining | protocols.io | dx.doi.org/10.17504/protocols.io.3byl4qoxrvo5/v1 | reuse | |
| **Protocol** | iPSC cultures | protocols.io | 10.17504/protocols.io.eq2ly523pvx9/v1 | new | |
| **Protocol** | Midbrain organoids differentiation | protocols.io | dx.doi.org/10.17504/protocols.io.rm7vzbnr4vx1/v1 | reuse | |
| **Protocol** | Midbrain neuronal culture | protocols.io | 10.17504/protocols.io.x54v9bzmml3e/v1 | new | |
| **Protocol** | NGN2 neuronal culture | protocols.io | 10.17504/protocols.io.j8nlk12ewg5r/v1 | new | |
| **Protocol** | Midbrain Astrocytes extraction and culture | protocols.io | dx.doi.org/10.17504/protocols.io.261ge364wl47/v2 | new | |
| **Protocol** | Co-cultures | protocols.io | 10.17504/protocols.io.rm7vze28xvx1/v1 | new | |
| **Protocol** | Conditioned Media Experiments | protocols.io | 10.17504/protocols.io.8epv55m1nv1b/v1 | new | |
| **Protocol** | Protein Extraction and Immunoblotting | protocols.io | dx.doi.org/10.17504/protocols.io.q26g7y428gwz/v1 | new | |
| **Protocol** | Lentiviral Production and Transduction | protocols.io | dx.doi.org/10.17504/protocols.io.6qpvr8ydolmk/v1 | reuse | |
| **Protocol** | Live-Cell imaging of α-synuclein internalization | protocols.io | 10.17504/protocols.io.261ge1jowv47/v1 | reuse | |
| **Protocol** | RNA Extraction and RT-qPCR | protocols.io | dx.doi.org/10.17504/protocols.io.3byl414krlo5/v1 | reuse | |
| **Protocol** | Immunofluorescence and Confocal Microscopy | protocols.io | dx.doi.org/10.17504/protocols.io.eq2lyqmemvx9/v1 | reuse | |
| **Protocol** | Effector Caspase assay | protocols.io | dx.doi.org/10.17504/protocols.io.6qpvrqb5blmk/v1 | reuse | |
| **Protocol** | LDH Cytotoxicity Assay | protocols.io | 10.17504/protocols.io.q26g776b3gwz/v1 | new | |
| **Protocol** | Lysosomal Function and Phagocytosis Assays | protocols.io | dx.doi.org/10.17504/protocols.io.14egn2xmyg5d/v1 | reuse | |
| **Protocol** | Cytokine Profiling | protocols.io | 10.17504/protocols.io.kxygx82eov8j/v1 | new | |
| **Protocol** | Polyamine Uptake Assay - FACS | protocols.io | 10.17504/protocols.io.q26g7mpq1gwz/v1 | new | |
| **Protocol** | Polyamine Uptake Assay - Confocal imaging | protocols.io | 10.17504/protocols.io.5qpvo9kd9v4o/v1 | new | |
| **Protocol** | Dotblot aggregated a-synuclein | protocols.io | dx.doi.org/10.17504/protocols.io.bp2l629x1gqe/v1 | new | |
| **Protocol** | Astrocytes Viral Transfection | protocols.io | dx.doi.org/10.17504/protocols.io.n2bvjbwxpgk5/v1 | new | |
| **Antibody** | α-synuclein | Abcam | #AB138501 | reuse | 1:300 staining, 1:1000 blots |
| **Antibody** | Actin | EMD Millipore | #S2532 | reuse | 1:2000 |
| **Antibody** | Aggregated-synuclein | Abcam | #AB209538 | reuse | 1:1000 |
| **Antibody** | ATP13A2 | Sigma | #A3361 | reuse | 1:1000 |
| **Antibody** | CD44 | Cell Signaling Technology | #3570S | reuse | 1:1000 |
| **Antibody** | FOXA2 | Abcam | #AB60721 | reuse | 1:500 |
| **Antibody** | GAPDH | Abcam | #ab9485 | reuse | 1:1000 |
| **Antibody** | GFAP | EMD millipore | #AB5804 | reuse | 1:300 |
| **Antibody** | GFAP | Dako | #AB_10013382 | reuse | 1:300 |
| **Antibody** | GIRK2 | Alomone labs | #APC-006 | reuse | 1:300 |
| **Antibody** | Histone 3 K9 dimethylation | Abcam | #AB_449854 | reuse | 1:300 |
| **Antibody** | Histone 3 K4 trimethylation | Abcam | #AB_306649 | reuse | 1:300 |
| **Antibody** | LAMP1 | Novus biotech | #NBP2-25183 | reuse | 1:300 |
| **Antibody** | MAP2 | BioLegend | #822501 | reuse | 1:500 |
| **Antibody** | S100A6 | Abcam | #ab181975 | reuse | 1:200 |
| **Antibody** | S100B | EMD millipore | #S2532 | reuse | 1:300 |
| **Antibody** | SOX9 | Abcam | #AB185966 | reuse | 1:1000 |
| **Antibody** | TH | Abcam | #ab112 | reuse | 1:300 |
| **Antibody** | TUJ1 | BioLegend | #801202 | reuse | 1:300 |
| **Chemical, peptide, or recombinant protein** | DAPI | Cayman Chemical | #13197 | reuse | 1:5000 |
| **Chemical, peptide, or recombinant protein** | Recombinant aSynuclein-HiLyte Fluor 488 labeled | Anaspec | #AS-55457 | reuse | |
| **Chemical, peptide, or recombinant protein** | CXCL1 | Peprotech | #275-GR-010 | reuse | |
| **Software/code** | FIJI Version 2.10.0 | National Institute of Health (NIH) | https://imagej.net/software/fiji/; RRID: SCR_002285 | reuse | |
| **Software/code** | Cell Profiler | Broad Institute | https://cellprofiler.org | reuse | |
| **Software/code** | GraphPad Prism Version 10.6.0 | | https://www.graphpad.com | reuse | |
| **Dataset** | Quantification data | Zenodo | 10.5281/zenodo.18704033 | new | |
